# SARS-CoV-2 infection leads to sustained testicular injury and functional impairments in K18 hACE2 mice

**DOI:** 10.1101/2023.10.31.565042

**Authors:** Stefanos Giannakopoulos, Monika A Ward, Jackson Bakse, Jin Pak, Vivek R Nerurkar, Michelle D Tallquist, Saguna Verma

**Affiliations:** Department of Cell and Molecular Biology, John A. Burns School of Medicine, University of Hawaii at Manoa, Honolulu, Hawaii, USA; Institute for Biogenesis Research, John A Burns School of Medicine, University of Hawaii at Manoa, Honolulu, Hawaii, USA; Department of Tropical Medicine, Medical Microbiology, and Pharmacology, John A. Burns School of Medicine, University of Hawaii at Manoa, Honolulu, Hawaii, USA; Center for Cardiovascular Research, John A. Burns School of Medicine, University of Hawaii at Manoa, Honolulu, Hawaii, USA

## Abstract

Compromised male reproductive health is one of the symptoms of long COVID with a decrease in male fertility markers including testosterone levels and sperm count for months in recovering patients. However, the long-term impact of SARS-CoV-2 infection on testicular injury and underlying mechanisms remains unknown. We previously demonstrated a disrupted tissue architecture with no evidence of virus replication in the testis during the acute stage of the disease in K18-hACE2 mice. Here, we systematically delineate the consequences of SARS-CoV-2 infection on the testis injury and function both during the acute stage of the disease and up to 4 weeks after infection in survivor K18-hACE2 mice. The gross morphological defects included sloughing of healthy spermatids and spermatocytes into the lumen, lack of lumen, and increase in apoptotic cells that sustained for at least 2 weeks after infection. Testis injury correlated with systemic and testicular inflammation, and infiltration of immune cells in the interstitial space and seminiferous tubules. Transcriptomic analysis identified dysregulation of key pathways of testicular immune homeostasis, spermatogenesis, and cell death at the symptomatic and short-term recovery stages. Further, a significant reduction in testosterone levels was associated with transient reduction in sperm count and mouse fertility. Most of the testicular impairments except testosterone levels were resolved within 4 weeks, which is almost one spermatogenesis cycle in mice. These findings provide much-needed mechanistic insights beyond our current understanding of testicular pathogenesis, suggesting that recovering COVID-19 patients should be closely monitored to rescue the pathophysiological effects on male reproductive health.

## Introduction

The ongoing coronavirus disease 2019 (COVID-19) pandemic caused by the severe acute respiratory syndrome coronavirus 2 (SARS-CoV-2) remains a major health concern even after new therapies and vaccinations that have diminished acute fatality rates. A high percentage of symptomatic and asymptomatic COVID-19 survivors develop post-acute sequelae of infection (PASC) or ‘long COVID’ (1–3). The incidence of long COVID is estimated at 10–30% of non-hospitalized cases, 50–70% of hospitalized cases, and 10–12% of vaccinated cases, with the highest percentage of diagnoses between the ages of 36 and 50 years (4, 5). Long COVID is a multisystemic condition comprising often severe symptoms including chronic kidney disease, cardiovascular dysfunction, and compromised male reproductive system that continues for weeks/months after recovery from infection of different SARS-CoV-2 variants (1, 4).

Strong clinical data report orchitis (pain and discomfort due to inflammation) as one of the symptoms during the acute stage of the disease in almost a quarter of infected men (4, 6). Postmortem analyses of testes from COVID-19 patients have revealed signs of mild to severe testicular pathology, including testicular swelling, tubular injury, germ cell and Leydig cell (LC) depletion, and leukocyte infiltration (7–10) that directly correlated with altered fertility parameters. The alterations in male fertility parameters like reduced sperm count, decreased testosterone levels, and dysregulated ratio of testosterone to luteinizing hormone (T/LH) have been reported in COVID-19 patients suggesting primary testicular failure with LC involvement (11, 12). The presence of SARS-CoV-2 RNA in the testis and semen is a rare event (13–15). However, a recent longitudinal cohort study indicated a significant increase in inflammatory mediators including IL-6 in the semen that persisted for >2 months after recovery and correlated with significant impairments in the sperm morphology, concentration, and total count (6). Notably, even patients with moderate COVID-19 symptoms display statistically significant impairment of sperm parameters including reduced concentration and motility for months after recovery (6, 16). Further, hypogonadism 3-9 months after recovery is now well established as one of the symptoms of a compromised male reproductive system in the long COVID patients (17, 18). However, the underlying mechanisms of testicular injury and impaired function are not clear.

The immune privileged environment of the testis is tightly governed by an elaborate communication network between different somatic resident cells, including Sertoli cells (SC), that nurse undifferentiated spermatogonial stem cells (SSC) and form the blood-testis barrier (BTB), and LC required for maintaining spermatogenesis and local immune homeostasis (19). Viruses can impact testis functions and male fertility by either dysregulating the hypothalamus-pituitary-gonadal (HPG) axis or by a direct infection of testicular cells (20). Since ACE2, the entry receptor of SARS-CoV-2, is highly expressed in testicular cells, we and others earlier speculated that SARS-CoV-2 can productively infect resident cells. However, our previous study using human 2-dimensional (2D) and 3-dimensional (3D) testicular culture systems provided evidence that SARS-CoV-2 does not establish productive infection in human testicular cells and that the testicular injury is not due to direct infection of SARS-CoV-2 but more likely is an indirect effect of systemic inflammation (21). Systemic cytokine storm during acute SARS-CoV-2 infection is characterized by the presence of elevated levels of pro-inflammatory cytokines including IL-1β, IL-6, TNF-α, GM-CSF, IFN-γ, MCP-1 and VEGF (22) and linked to multiple-organ damage and immune dysfunction during both acute stage of disease and in long COVID patients (23, 24). Our data obtained with the K18-hACE2 transgenic mice demonstrated gross pathology in the testis during the acute stage of the disease (21) similar to testicular injury observed in post-mortem COVID patients, thus supporting the use of these mice to assess the long-term impact of SARS-CoV-2 infection on testes.

The extent of testicular injury post-recovery from COVID-19 and the mechanisms underlying functional impairments are not yet clear. In this study, we systematically delineate the impact of SARS-CoV-2 infection on different aspects of testicular injury and function markers in mice, both during the acute stage of the disease and up to 30 days post-infection, which is almost one spermatogenesis cycle in mice. The data provides direct evidence of sustained testicular injury, inflammation, and infiltration of immune cells during recovery. Transcriptomics analysis identifies dysregulation of key pathways associated with inflammation, spermatogenesis, steroidogenesis, and cell death that correlates with a significant reduction in testosterone levels. Further, reduced sperm counts in recovering mice were associated with a transient decline in the fertility of mice. However, most of the injury markers and defects resolved within 4 weeks of infection. Collectively, the data provide much-needed mechanistic insights into the effects of SARS-CoV-2 infection on testis that will inform the efforts to monitor and rescue pathophysiological effects of COVID-19 on male reproductive health.

## Results

### Severe morphological alterations in the testis are observed during the acute stage of the disease and the recovery stage in SARS-CoV-2-infected hACE2 mice

We previously reported that SARS-CoV-2 infection in K18-hACE2 mice leads to gross histopathological changes in the testis during the acute stage of the disease in the absence of any active virus replication in the testis (21). To further understand the long-term implications of the infection on the injury and to quantify different types of defects at different stages of the disease, we monitored survivor mice for up to 30 days post-infection (D30). The SARS-CoV-2 infection of K18-hACE2 mice resulted in 80% mortality (Fig. S1A). The survivor mice lost body weight drastically during the acute stage of the disease (peak at D5) and their recovered body weight remained lower than the mock-infected group (Fig. S1B). As expected, the virus was cleared from the lungs by D8 (Fig. S1C and D) and there was no infectious virus detected in the testis as shown in our previous study (21) and Fig. S1D.

To gain more insights into the testicular alterations during the extended timeframe post-infection, we characterized testicular abnormalities using testis sections during the pre-symptomatic stage when virus replication is well established in lungs (D3), peak symptomatic stage (D5), short-term recovery stage when the virus is cleared from lungs (D8 and D14), and long-term recovery stage (D30), which is almost one spermatogenesis cycle in mice. Testes from control (mock-infected) mice showed normal spermatogenesis, with the germ cell types and organization as expected (Fig. 1Ai). When testes from infected mice were assessed, the support cells in the interstitial space (LC) and seminiferous epithelium looked normal at all examined time points. At D3 (Fig. 1Aii) the testes resembled those of control mice. At D5, D8, and D14 interstitial edema was observed (Fig. 1Aiii-iv) and many of the seminiferous tubules were classified as abnormal, with predominant defects including lack of clear lumen and separation of germ cell layers from the basal membrane (Fig. 1Aiii) and germ cells sloughing into the lumen (Fig. 1Aiv-v). The abnormalities of individual germ cells were also observed, with some cells presenting with degenerating nuclei (Fig. 1Aiii) and undergoing apoptosis (Fig. 1Aiv-v). At D30, the appearance of the germ cell and seminiferous epithelium was normal (Fig. 1Avi). Quantification of tubular and germ cell defects showed no effects at D3, severe abnormalities at D5, partial retention of defects at D8-14, and complete recovery at D30 (Fig. 1B). Overall, these data suggest that SARS-CoV-2-induced testicular defects during the acute stage of the disease persist for at least 2 weeks post-infection but are completely resolved by D30 in the K18-hACE2 mouse model.

**Figure 1.**
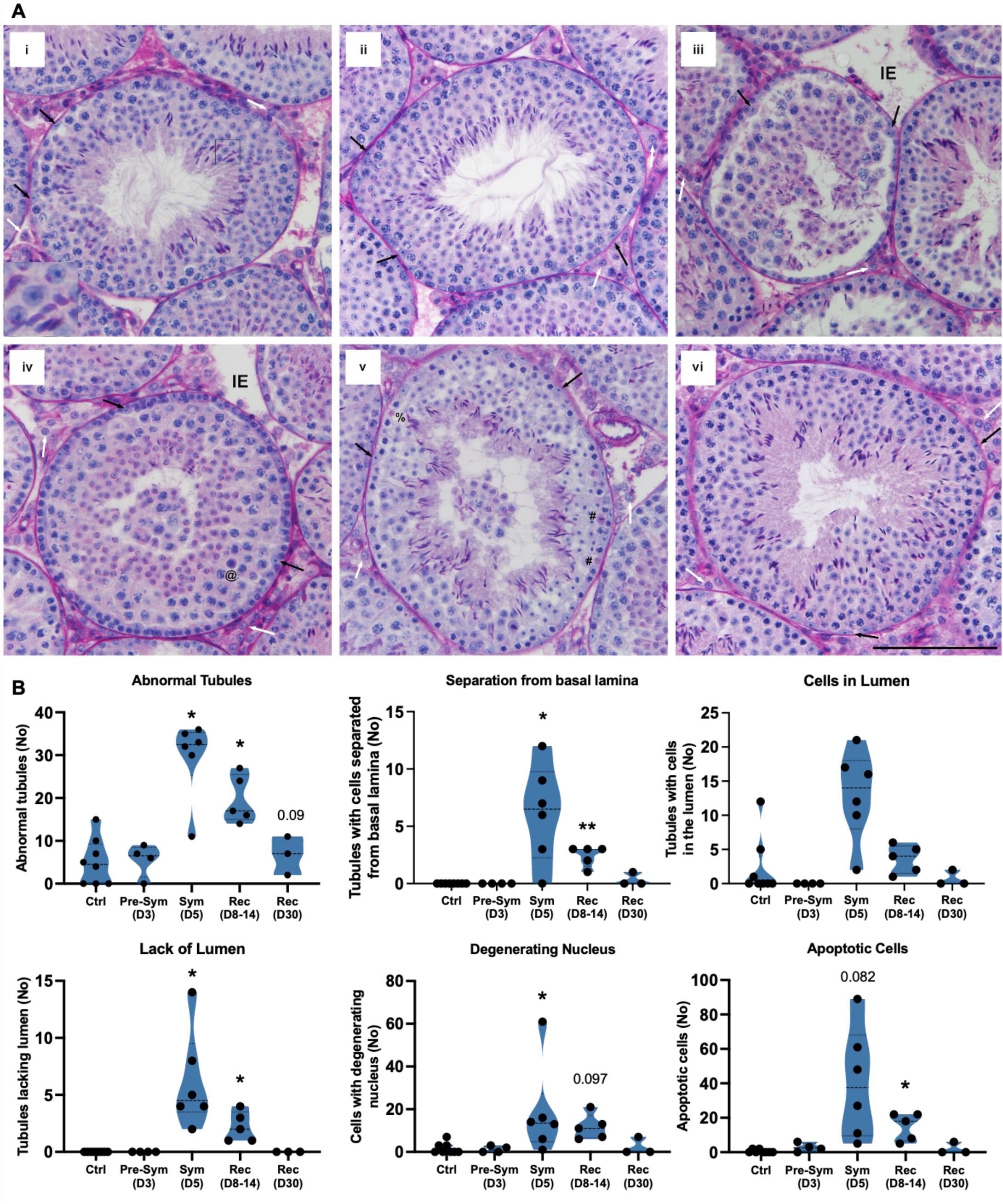
SARS-CoV-2-infection induces severe morphological defects in the testis during the acute stage of the disease that persist for short-term during recovery in hACE2 mice. Seminiferous tubule organization and germ cell normalcy were assessed in testes from K18-hACE2 mice sacrificed after mock (i, Ctrl) or SARS-CoV-2 (ii-vi, infected) infection. The days post-infection (D) are represented as pre-symptomatic (D3), symptomatic (D5), recovering (D8-14), and recovered (D30). **(A)** Exemplary seminiferous tubules of PAS-H-stained testis sections. Control seminiferous tubules (i) show normal spermatogenesis and the presence of the expected germ cell types and organization, with healthy support cells in the interstitial space (Leydig cells, white arrows) and in the seminiferous epithelium (Sertoli cells, black arrows). Inset shows normal round and elongating spermatids. At D3 (ii), seminiferous tubule organization and germ cell health remained similar to that of control males. Beginning at D5 (iii) and through D8 (iv) and D14 (v) various testicular abnormalities were observed, including interstitial edema (IE), occurrence of apoptotic germ cells (@ in iv), apoptotic meiotic cells (# in v), and cells with degenerating nuclei (% in v), as well as tubules with a lack of clear lumen, a separation of germ cell layer from the basal membrane (iii), and with the sloughing of healthy spermatids and spermatocytes into the lumen (iv, v). Leydig and Sertoli cells appeared healthy following SARS-CoV-2 infection (ii-vi; white and black arrows, respectively). Germ cell and seminiferous epithelium were normal at D30 (vi). The stages of seminiferous epithelium are: II-IV (vi), V-VI (i, ii) VII-VIII (iii, iv), X-XII (v). Inset, 3x magnification. Scale, 100 µm. **(B)** Quantification of seminiferous tubule organizational and germ cell defects. For each male, 50 tubules were examined and classified as normal or abnormal in regard to their organizational and germ cell features. Each data point corresponds to a different mouse. Statistical significance (t-test, Ctrl vs. Infected); *, p<0.05. The differences approaching significance (p=0.05-0.1) are shown directly in the graph.

### RNA-seq analysis of the testis post-SARS-CoV-2 infection reveals alterations in key pathways associated with testis function

To gain insights into the impact of SARS-CoV-2 on the testes at the molecular level, we examined the transcriptomic signatures in the testes from infected K18-hACE2 mice by RNA-seq analysis. Gene signatures were analyzed at D3, D5, D8, D14 and D30. Total reads, aligned reads, percent alignment, average read length, percent GC content, Phred score, and Q30 are shown in Fig. S2. The transcriptome profiling and comparison with mock-infected samples revealed a total of 1,865 non-redundant differentially expressed genes (DEGs) across all time points (p<0.05). As shown in the Volcano plots and the Venn diagram (Fig. 2A-B) the highest number of timepoint-unique DEGs were observed at D8 (691). The highest number of upregulated genes was observed at D30 (386 upregulated vs, 96 downregulated) unlike other time points. Notably, a high number of dysregulated genes were common between many time points indicating that infection results in both common and distinct gene signatures at each time point. Principal component analysis (Fig. 2C) showed that the control testis and the infected samples were distinctly separated, with most of the infected samples grouped together. Gene ontology (GO) overrepresentation analysis (ORA) of DEGs at D3 and D5 revealed the downregulation of genes associated with cell adhesion and extracellular structure reorganization (Fig. S3A), while genes associated with cytokine and chemokine secretion and lymphocyte differentiation pathways were upregulated at D8 and D14 (Fig. S3B). The most prominently upregulated pathway at D30 was the response to wounding and wound healing while the downregulated pathways included tissue homeostasis (Fig. S3C).

**Figure 2.**
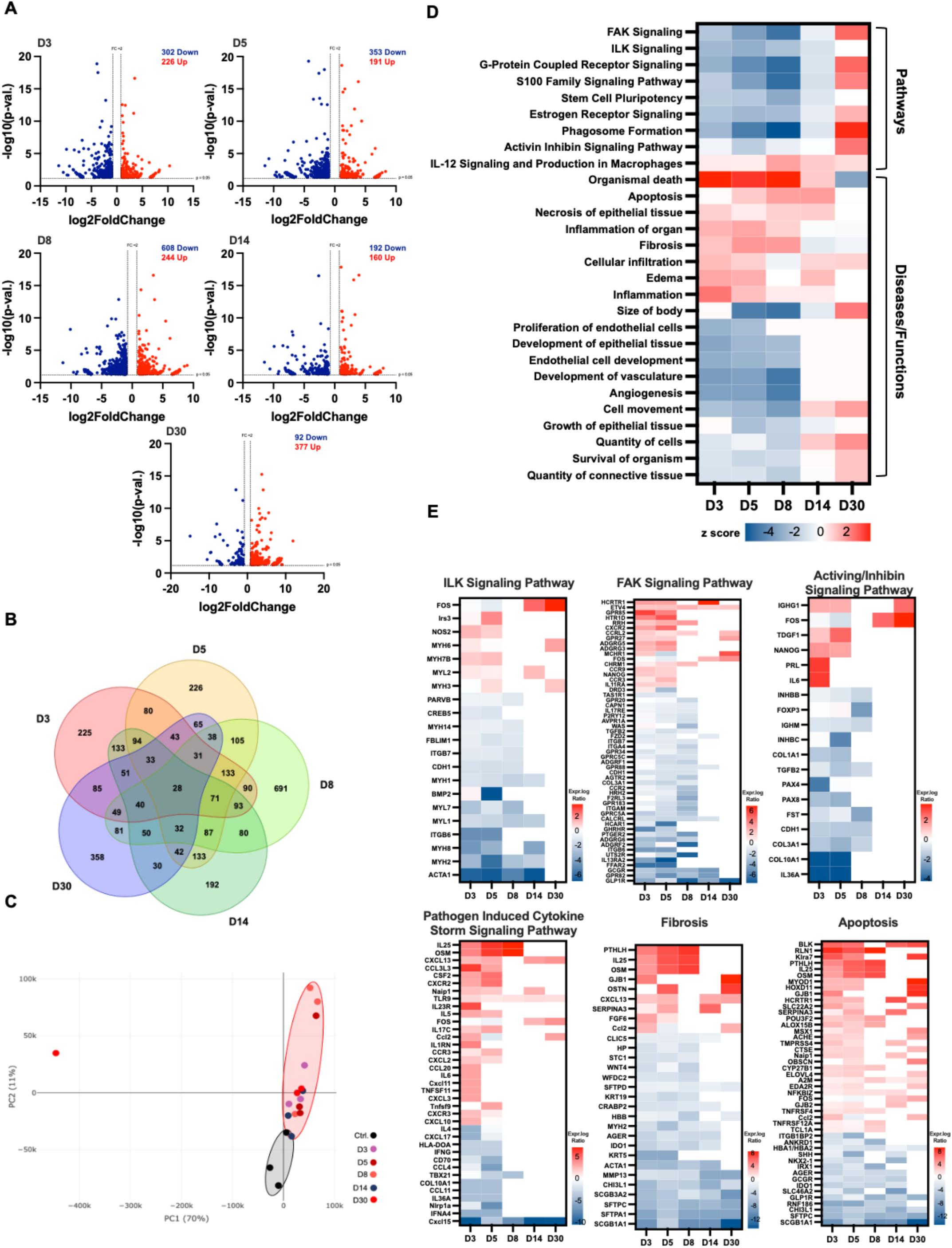
RNA-seq analysis of whole testis identifies dysregulation of distinct transcriptional pathways at different stages of SARS-CoV-2 infection. **(A)** Volcano plots of differentially expressed genes (DEGs) (n, 3 per time point) isolated at D3, D5, D8, D14, and D30, dotted lines represent cutoff: p< 0.05 and log2fc > |1|, upregulated, red; and downregulated, blue. **(B)** Venn diagram of total DEGs at D3 (red), D5 (yellow), D8 (light green), D14 (dark green), and D30 (blue). **(C)** Principal component analysis (PCA) performed using DEGs (p <0.05 and log2fc > |1|) and normalized TPM (controls, black circle; infected, red circle) **(D)** Heatmap of top dysregulated canonical pathways and select diseases and functions revealed by Ingenuity Pathway Analysis (IPA), filtered to remove erroneous results (upregulated, red; downregulated, blue) at D3, 5, 8,14, and 30. **(E)** Expression log ratio of upregulated (red) and downregulated (blue) genes associated with ILK, FAK, and activin/inhibin signaling pathways and pathogen-induced cytokine storm, fibrosis, and apoptosis signaling pathways as determined by IPA in each dataset (p< 0.05; log2fc > |1|) at indicated time points.

The DEGs based on their significance (p< 0.05) and log2fold change (cutoff >|1|) were further processed for pathway enrichment using Ingenuity Pathway Analysis (IPA) to identify the top functional networks modulated in the testis at different time points following SARS-CoV-2 infection (Fig. 2D-E). At early time points (D3-8), there were more downregulated pathways that included the focal adhesion kinase (FAK) signaling pathway and the integrin-linked kinase (ILK) signaling pathway. These pathways play an important role in tight junction formation and cell-cell adhesion (25, 26), indicating the potential disruption of the blood-testis barrier (BTB) with a peak at D8. The signaling pathways associated with stem cell pluripotency, estrogen receptor, and phagosome formation were downregulated at D3-D14, while the activin-inhibin pathway was transiently downregulated only at D5. The IPA analysis of select diseases and functions revealed significant upregulation of organismal death, cellular infiltration, apoptosis, necrosis, inflammation of organs, and fibrosis throughout the time points that were resolved only by D30. The IL-12 signaling was upregulated at D8, D14, and D30, indicating potential activation of testicular immune cells. Notably, other key diseases/functions, such as cell proliferation, development, angiogenesis, cell movement, and survival were also downregulated starting at early time points.

Individual DEGs of select pathways/functions are shown in Fig. 2E. ILK signaling pathway includes ACTA1, a skeletal muscle α-actin gene that plays a role in muscle contraction and cytoskeleton organization (27), that was highly downregulated at all the time points besides D30. INHBB and TGFB2, are both involved in activin/inhibin signaling pathway and were significantly downregulated only at early time points. Transcript of Oncostatin-M (OSM), which is a multi-functional cytokine that promotes inflammation (28) and negatively regulates Leydig cell differentiation (29) was significantly upregulated and was part of pathogen-induced cytokine storm, fibrosis, and apoptosis pathways. Additional cytokines and chemokines induced in these pathways included IL-25, CXCL13, CSF2, CXCL2, CCL2, Naip1, and members of the TNF family (TNFSF11 and TNFSF9). Furthermore, consistent with the absence of active viral replication, the interferon genes such as IFNG and IFNA4 and ISGs were downregulated or not induced at any time points. However, CXCL15, which has been shown to suppress proliferation and expansion of multi-potential, erythroid, granulocyte, and macrophage progenitors (30), was significantly downregulated at all time points. Taken together, these data indicate that pathways associated with testicular immune homeostasis were significantly affected in the initial stages of infection and in short-term recovery stage, however, genes associated with apoptosis remained dysfunctional till D30.

### IPA upstream regulator analysis reveals significant alterations in fertility markers

We then employed IPA’s predictive upstream regulator analysis to predict activation or inhibition of transcription regulators that may lead to differential expression of genes observed in Fig 2. Upstream regulators related to fertility were downregulated at D3, D5 and D8 (Fig. 3A) while regulators of organismal cell death were upregulated at these time points. The androgen receptor (AR), an important steroid hormone receptor that influences the sustenance of spermatogenesis pathways, was amongst the top downregulated upstream molecules, which was identified to be low even at D30. Angiotensinogen (AGT), a member of the Renin/ACE/ AngII/AT1R/AT2R axis that plays an important role in testicular growth/differentiation and is mainly produced by Leydig cells (31), was also identified to be downregulated in the early timepoints, suggesting Leydig cell dysfunction in the acute stage of the disease (Fig. 3B). GATA1, GATA2 and GATA6 are important transcription factors in steroidogenesis and testosterone production (32–34) and they were all downregulated in the early timepoints with GATA6 showing downregulation even at D30 (Fig. 3B). Therefore, we next validated the association of the expression of GATA6 and AR with the downstream genes targeted by these regulators. Steroidogenic genes SFTPC, SFTPA1, and SCGB1A1 regulated by GATA6 were significantly downregulated at all the time points with the latter having a fold change of -14.9 even at D30 (Fig. 3C). CYP2J13, TNNT3, and PVALB transcripts targeted by the androgen receptor (AR) were also downregulated at all time points except D30 (Fig. 3D). We also observed that VEGF, a growth factor associated with angiogenesis, and TGFB1, a key player in the immune suppressive environment of the testis, were predicted to be downregulated in the early time points (Fig. 3B). In contrast, mediators of apoptosis (THZ1, a CDK7 inhibitor, (35), fibrosis (SIX1), and inflammation (TNFSF12) were upregulated in the early time points. These data collectively suggest that important regulators of spermatogenesis, steroidogenesis, and immune homeostasis are adversely affected by SARS-CoV-2 infection.

**Figure 3.**
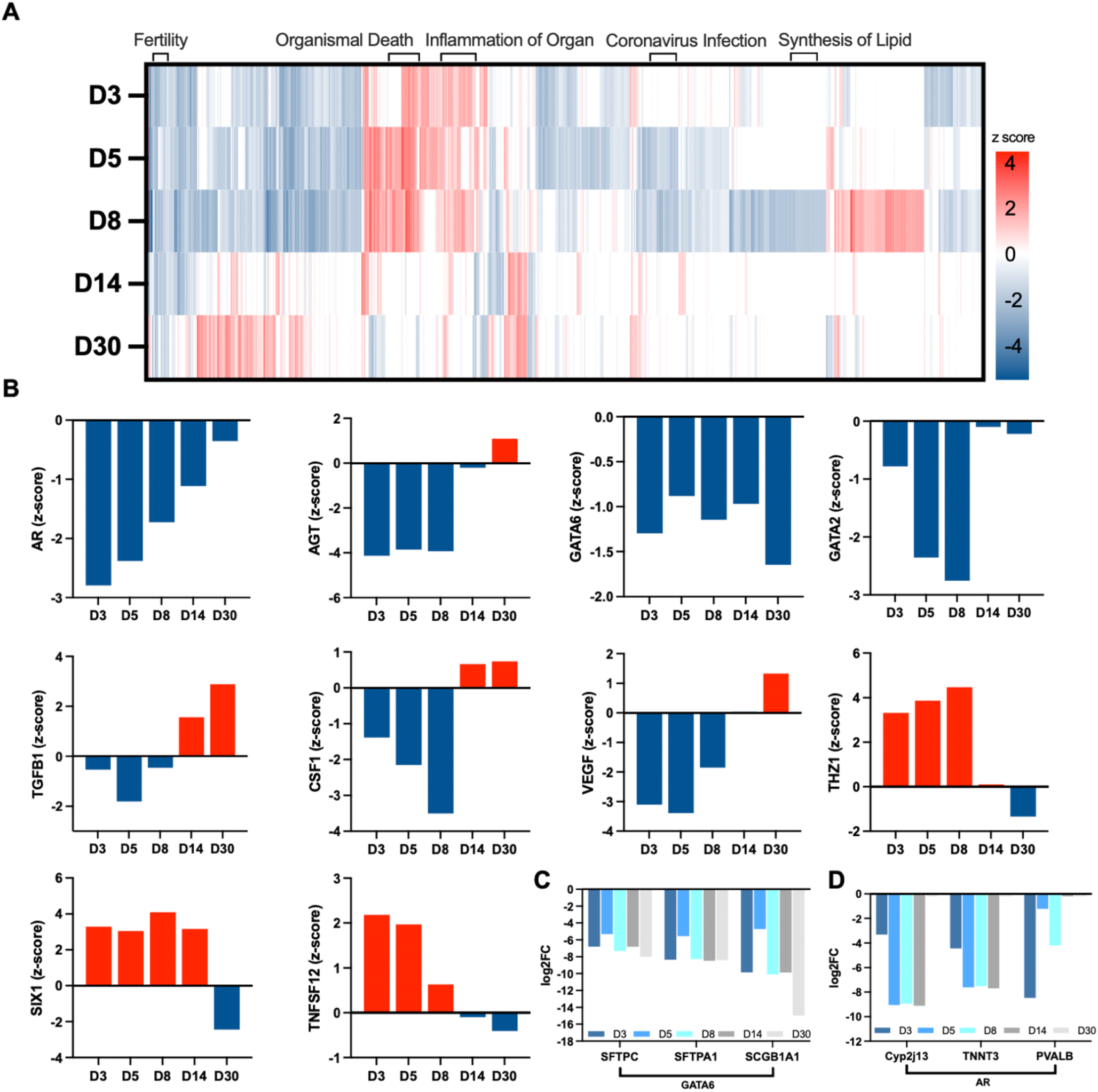
Ingenuity pathway analysis of upstream regulators reveals significant alterations in fertility and cell death markers following SARS-CoV-2 infection of K18hACE2 mice. **(A)** Heatmap of hierarchical clustering of global upstream regulators modulated during SARS-CoV-2 infection identified by IPA at each time point (n, 3 per time point). Brackets distinguish clusters of upstream regulators associated with fertility, organismal death, inflammation of organ, coronavirus infection and lipid synthesis (up, red; and down, blue). **(B)** z-score of select upstream regulators associated with fertility at indicated time points. The fold change of genes targeted by **(C)** GATA6 and **(D)** AR at indicated time points in testis from SARS-CoV-2 infected mice and expressed as log2fc. (p< 0.05; log2fc > |1|; n, 3 per group).

### Increased levels of pro-inflammatory cytokines and chemokines in the testis correlate with systemic inflammation

To validate the dysregulation of inflammatory pathways seen in our RNA-Seq data and to understand the correlation between systemic and testicular inflammation with testicular injury, we measured a panel of cytokines and chemokines specific for COVID-19 in the serum and lysates from lung, testis, and heart. Heatmap of relative protein levels of all 27 cytokines and chemokines with respect to controls is shown in Fig. 4A. As expected, levels of G-CSF, GM-CSF, IL-1α, IL-1β, IL-6, IL-15, CXCL10, CXCL1, CCL2, CCL3, CCL4, TNF-α, and IFN-γ increased significantly in lungs, and serum. Following what we have seen in serum, key inflammatory molecules including G-CSF, IL-1β, IL-6, CXCL1, CCL3, and TNF-α were also increased in the testis at D3-D14. The IL-6 and TNF-α levels in the testis remained significantly higher even at D30 than in the controls suggesting that an increase of specific testicular cytokines may take longer to completely resolve. As expected, IFN-γ levels that were high in lung and serum and indicative of active virus replication were not increased in the testis at any time point. Fig 4B compares the absolute levels of IL-1β, IL-6, CXCL1, and TNF-α well-established markers of SARS-CoV-2-induced cytokine storm in lung, serum, and testis. To examine the correlation between the cytokines in the serum (Fig. 4C left panel) and testis (Fig. 4C right panel) with gross morphological alterations observed in Fig. 1B, we performed Pearson correlation analysis. The correlation between different testicular abnormalities (basal lamina separation, lack of lumen, and cells in lumen) with testicular cytokines IL-1β, IL-6, and TNF-α (Fig. 4C right panel) was much stronger than with serum cytokines. The correlation values were weaker between the levels of three chemokines tested and testicular injury markers except for CXCL1 (Fig. 4C right panel). Collectively, this data highlights that the testicular immune environment is compromised for weeks after infection and shifts towards a robust proinflammatory response during the acute and short-term recovery stage of the disease.

**Figure 4.**
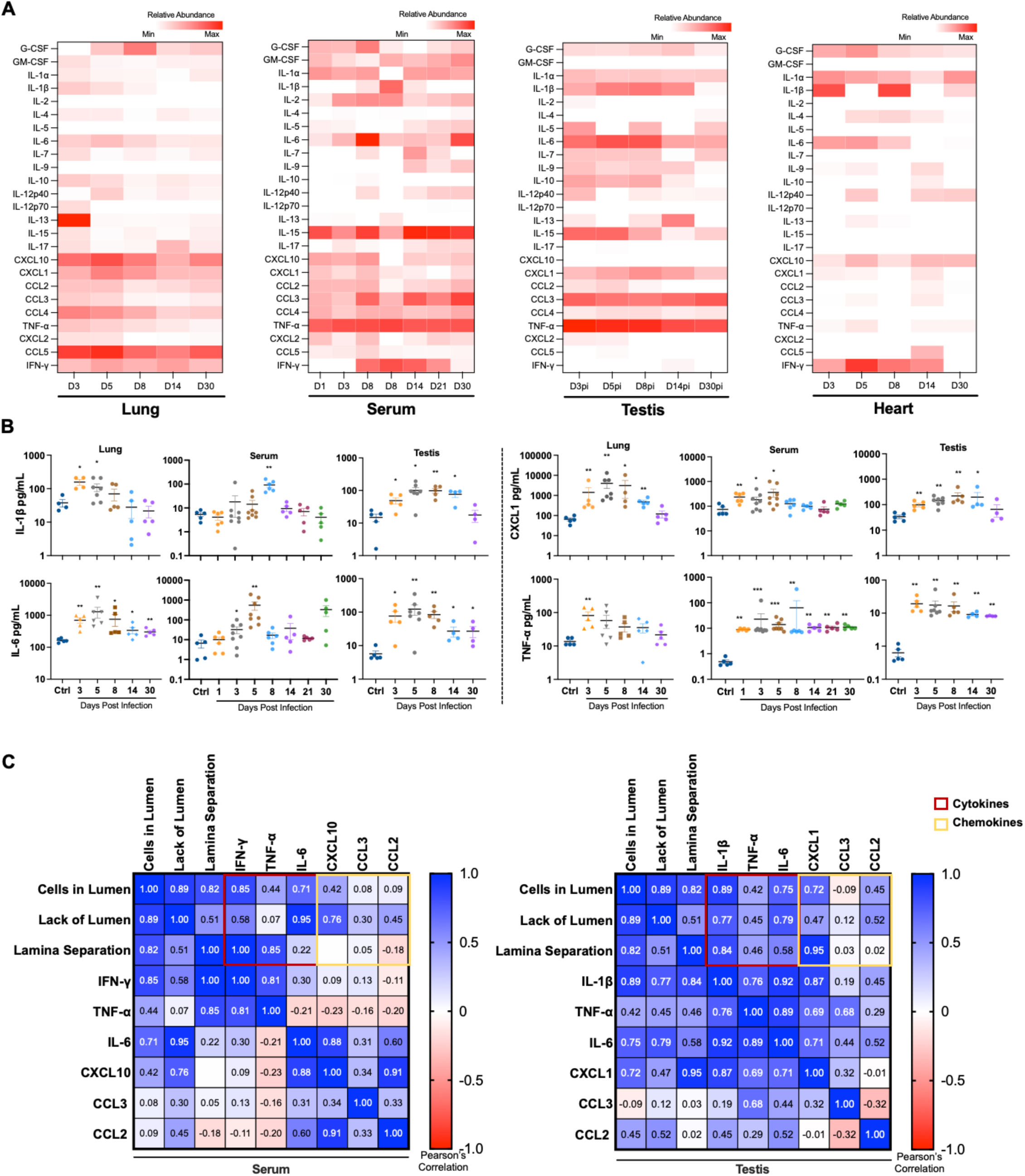
Increased levels of pro-inflammatory cytokines and chemokines in the testis following SARS-CoV-2 infection correlate with systemic inflammation. **(A)** Heatmap comparison of a panel of COVID-19 specific cytokine and chemokines measured using LUMINEX assay with the tissue homogenates (lung, testis, and heart) and serum of K18-hACE2 mice at indicated time points post-infection (n, ≥ 4 mice per group). Data were normalized to the control, log10 transformed, and expressed as relative abundance (red, increased; blue, decreased; and white represents unchanged with respect to uninfected controls). **(B)** Representative levels of IL-1β, IL-6, CXCL1 and TNF-α expressed as pg/mL. Individual data points represent different mice. The statistical significance (* p<0.05, **p<0.01, ***< p<0.001) was determined using Mann-Whitney test for all analyses **(C)** Pearson’s correlation coefficient analysis of three testicular defects shown in Fig. 1 and key cytokines (red box) and chemokines (yellow box) in serum (left) and testis (right) at D3, D5, and D8.

### Apoptotic cell death observed in the testis from infected mice correlates with infiltration of immune cells and loss of BTB proteins

To address the consequence of the pro-inflammatory response in the testis, we assessed cell death using Terminal deoxynucleotidyl transferase dUTP nick end labeling (TUNEL) assay at different time points (Fig. 5A). TUNEL staining of testis from control mice was comparable to that of D3 and revealed very low to no staining in either the seminiferous tubule or the interstitial space. At D5, a significant increase in TUNEL-positive cells was observed, primarily in the cells within the seminiferous tubules; some interstitial cells were also positive for TUNEL. At D8, TUNEL-positive cells peaked and were observed in both cells comprising seminiferous epithelium and those sloughed into the lumen. TUNEL-positive cells remained significantly more abundant when compared to testis from control mice even at D14 indicating that apoptotic cell death persists for at least 2 weeks after infection and after ∼7 days after clearance of SARS-CoV-2 from lungs. By D30, the incidence of TUNEL-positive cells reduced dramatically, although they were still more frequently observed than in control or D3 testis indicating that the testicular injury had not resolved completely (Fig. 5C). These TUNEL assay data correlated with the increase in apoptotic and organismal death pathways observed in our RNA-Seq data (Fig. 2D). The expression of BTB marker ZO-1 was found to reduced markedly at D5 and D8 (Fig. 5B and D) suggesting that testicular inflammation transiently affects BTB integrity. To gain insights into the transcript expression of other BTB-associated proteins, we interrogated other genes of gap junction and tight junction proteins families in our RNA Seq data (Fig. 5E). Many gap junction and tight junction genes specifically in the claudin family were downregulated at both D5 and D8 with some having more than -4 log2fc than the control.

**Figure 5.**
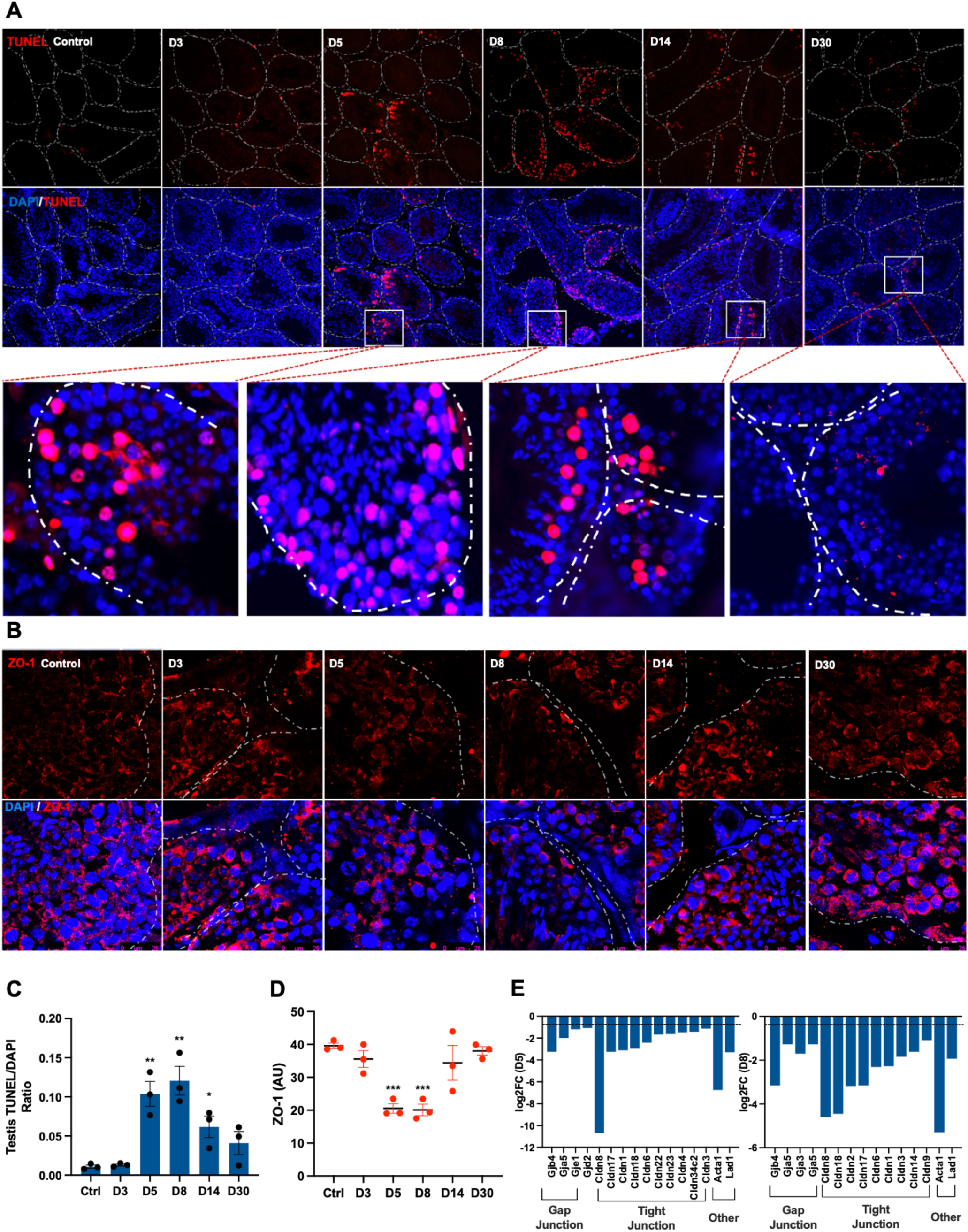
SARS-CoV-2 infection results in sustained apoptotic cell death in the testis of K18 hACE2 mice. **(A)** Representative images of TUNEL staining (red) and **(B)** ZO-1 staining (red) in the sections from OCT embedded testis at D3, 5, 8, 14 and 30 (n, 3 mice per time point). Nuclei were visualized using DAPI stain (blue) and dotted lines represent the shape of individual seminiferous tubules. ImageJ analysis of **(C)** TUNEL staining normalized with respective DAPI positive cells and **(D)** ZO-1 using ImageJ. Each data point is the average of counts from 3 different fields from each testis section (n, 3 mice/time point). **(E)** log2fc of gap junction, tight junction, and filament molecules at D5 (left) and D8 (right). Dotted line represents cutoff (log2fc > |1|, p <0.05). Significance (*p<0.05, ** p<0.01, *** p<0.001) was determined using a student’s t-test.

Increased pro-inflammatory cytokines and chemotactic factors such as CXCL1, and CCL3 as seen in Fig. 4A are directly associated with increased tissue immune cell infiltration. Therefore, we next examined lymphocyte infiltration in the testis, using well-established markers of inflammatory monocytes/macrophages. The staining for the CD68, a marker of monocytes/macrophages (Fig. 6A) was comparable between controls and D3 testis, however at D5 there was a significant increase of CD68-positive cells both in the interstitial space and inside the seminiferous tubules that remained elevated at D8. Similarly, staining of CD11b, a marker of activated myeloid cells revealed a high number of positive cells at D5 and D8 (Fig. 6B). Interestingly, at D8, CD11b+ cells were detected both inside the seminiferous tubules and in the interstitial space indirectly indicating compromised BTB. The incidence of both CD68+ and CD11b+ cells was comparable to the control group at D14 and D30. Collectively these data suggest that SARS-CoV-2 infection leads to sustained apoptosis in the testis and infiltration of immune cells is one of the transient effects of the SARS-CoV-2-associated testicular pathogenesis that correlates with loss of BTB proteins.

**Figure 6.**
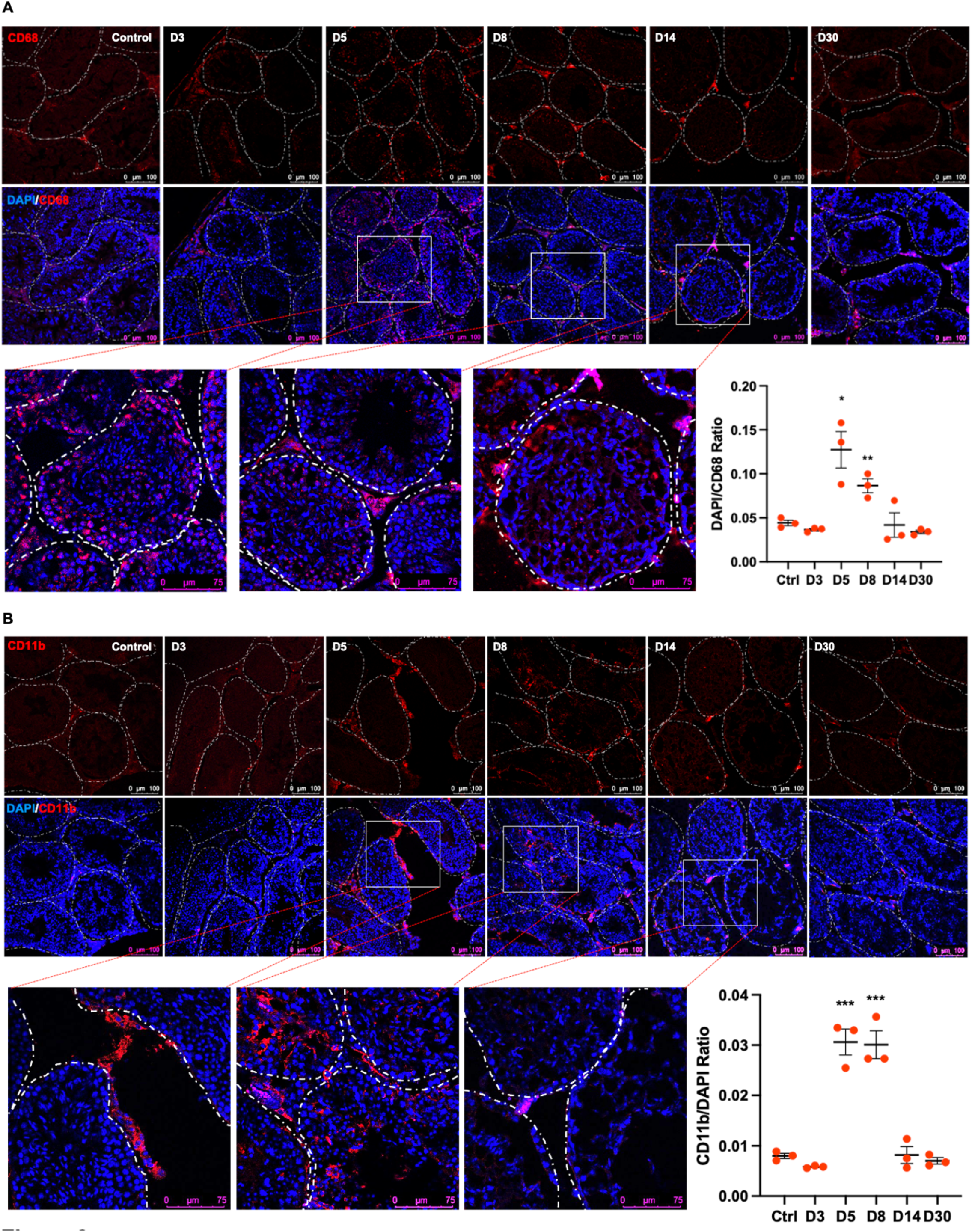
SARS-CoV-2 infection leads to increased infiltration of CD68+ and CD11b+ cells in the testis. Representative images of testis sections at D3, 5, 8, 14, and 30 with nuclei stained with DAPI (blue) and infiltrating leukocytes visualized using **(A)** CD68 (red), and **(B)** CD11b (red) antibodies. Dotted lines outline the shape of individual seminiferous tubules and ImageJ analysis of CD68 and Cd11b staining normalized to DAPI-positive cells was done using particle analyzer tool. Each data point is the average of counts from 3 different fields from each testis section (n, 3 mice/time point). Images were taken using Leica SP3, DMI8 confocal microscope. Significance (*p<0.05, **p<0.01, ***p<0.001) was determined using student’s t-test.

### Testicular function is compromised in SARS-CoV-2 infected hACE2 mice

The observation of gross testicular abnormalities, inflammation, and apoptotic cell death led us to another important question-what is the effect of all these alterations on testicular function measured as its ability to produce key hormones? We therefore assessed the levels of testosterone, follicular stimulating hormone (FSH), inhibin B (INHB), and luteinizing hormone (LH). Serum testosterone levels declined rapidly by D5 in a subset of mice and by D8 the decrease was significant and reached the lowest levels in all mice by D14 (Fig. 7A). Even though the testosterone levels slightly increased at D21 and D30 (Fig. 7A), they remained significantly lower than that of the control and D3 mice. The serum FSH levels were significantly elevated compared to controls at D8 and D14 (Fig. 7B). The INHB levels declined only at D5, and the LH levels remained comparable to the control group at all time points (Fig. 7C-D). To further understand the trend of these hormones over time, we performed linear regression analysis (Fig. 7E-G). Testosterone and inhibin B showed a negative correlation while FSH positively correlated (Pearson r=-0.9040, r=-0.6634, r=0.9795 respectively), with the disease progression (D1 to D8). The ratios of INHB/FSH, and Test/FSH, commonly used as a better indicator of spermatogenesis than either one alone (36, 37), showed a negative correlation from D1 to D8. A Pearson correlation matrix (Fig. 7J) demonstrated a negative correlation between FSH and INHB (r=-0.16), FSH and testosterone (r=-0.83), FSH and INHB/FSH ratio (r=-0.95), INHB and FSH/Test ratio (r=-0.12), testosterone and FSH/testosterone ratio (r =-0.81), and Inhibin B/FSH ratio and FSH/testosterone ratio (r=-0.93) further highlighting that SARS-CoV-2 infection leads to the imbalance of the key hormones involved in HPG axis and spermatogenesis in these mice.

**Figure 7.**
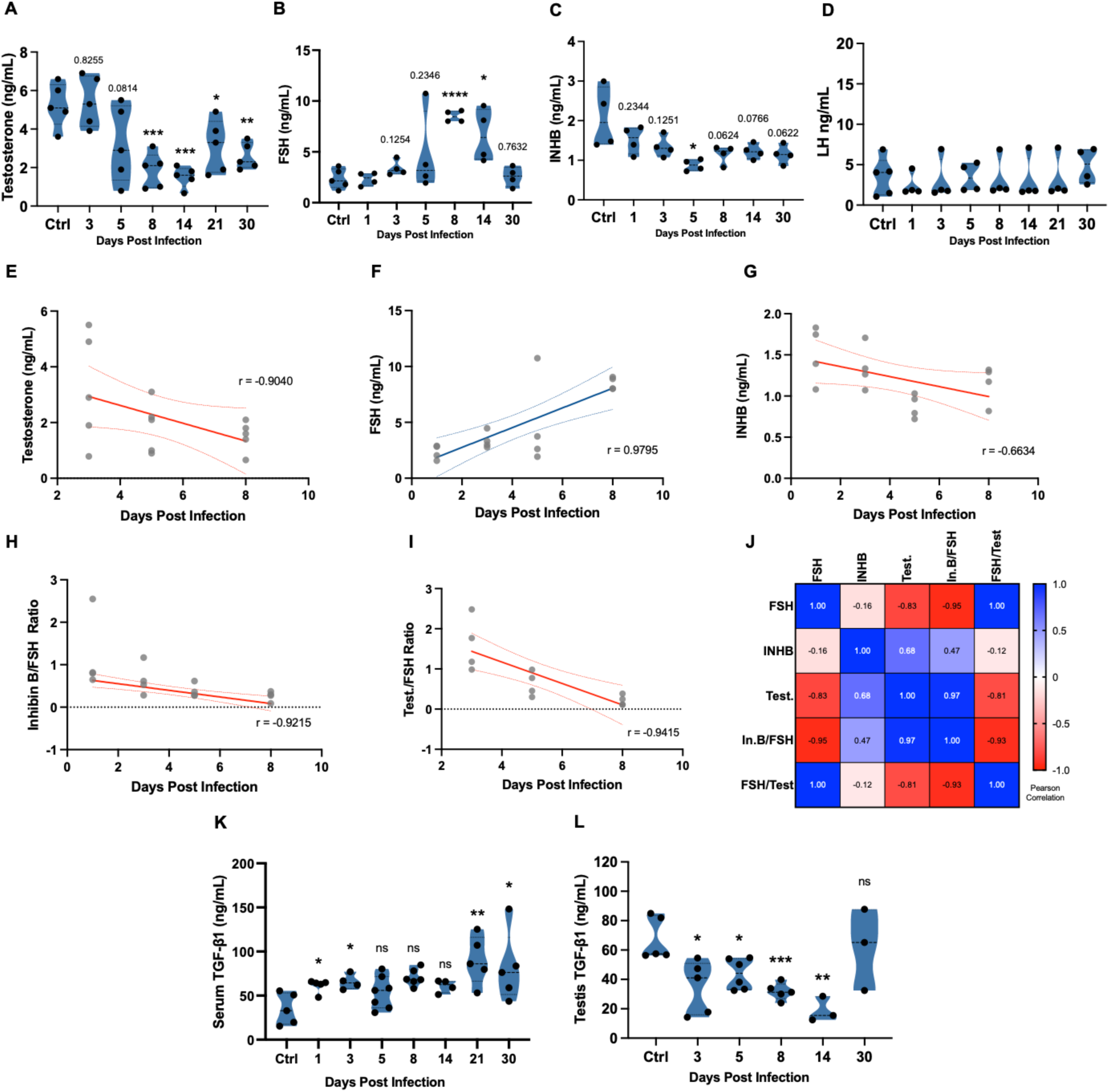
Testicular function is compromised in SARS-CoV-2-infected hACE2 mice. **(A)** Serum levels of testosterone, **(B)** follicle-stimulating hormone (FSH), **(C)** inhibin B (INHB), and **(D)** luteinizing hormone (LH) were measured using ELISA (n, at least 4 mice per group) at indicated time points and expressed as ng/mL serum. **(E-G)** Linear regression analysis of serum testosterone, FSH and INHB at D1, 3, 5, and 8 **(H-I)** Linear regression analysis of INHB/FSH and FSH/testosterone ratios performed at indicated time points. **(J)** Pearson coefficient correlation analysis of serum levels of testosterone and ratios of INHB/FSH and FSH/testosterone performed at D3, 5, and 8. TGF-β levels in serum **(K)** and testis homogenate **(L)** were measured using ELISA and expressed as ng/mL at indicated time points post-infection. Significance (*p<0.05, **p<0.01, ***p<0.001, ****p<0.0001) was determined using student’s t-test.

Considering the important role TGF-β plays in maintaining the immunosuppressive environment, we next measured testicular and serum levels of TGF-β1 (Fig. 7K and L). While serum TGF-β1 levels increased after infection and resembling what has been shown in human COVID-19 patients (38), we observed a gradual decline in testicular TGF-β1 levels with the lowest levels seen at D14, following which they recovered by D30.

Finally, a preliminary analysis of epidydimal sperm count and fertility was performed to test the downstream effect of the alterations in the testis function markers. There was a decline in the sperm count at D5, that became significant at D8 (∼50% reduction, Fig. 8A) with a trend of recovery at D15. To evaluate if decreased sperm count affected the ability to reproduce, the fertility was tested for a subset of the recovered mice. Three males, who were proven fertile before the infection and who survived the acute stage of infection, were paired with 2 females each at D10 for 6 days. Females paired with only 2 out of the 3 tested males became pregnant. The relationship between the sperm counts and the number of embryos in the females paired with each recovered male in Fig. 8B shows that the females paired with the male with the lowest sperm count did not get pregnant. Collectively, these data provide evidence that following SARS-CoV-2 infection, several markers of the testis injury and function are altered both during the acute and short-term recovery stage of the disease and our pilot data support that this can affect the fertility of survivor mice.

**Figure 8.**
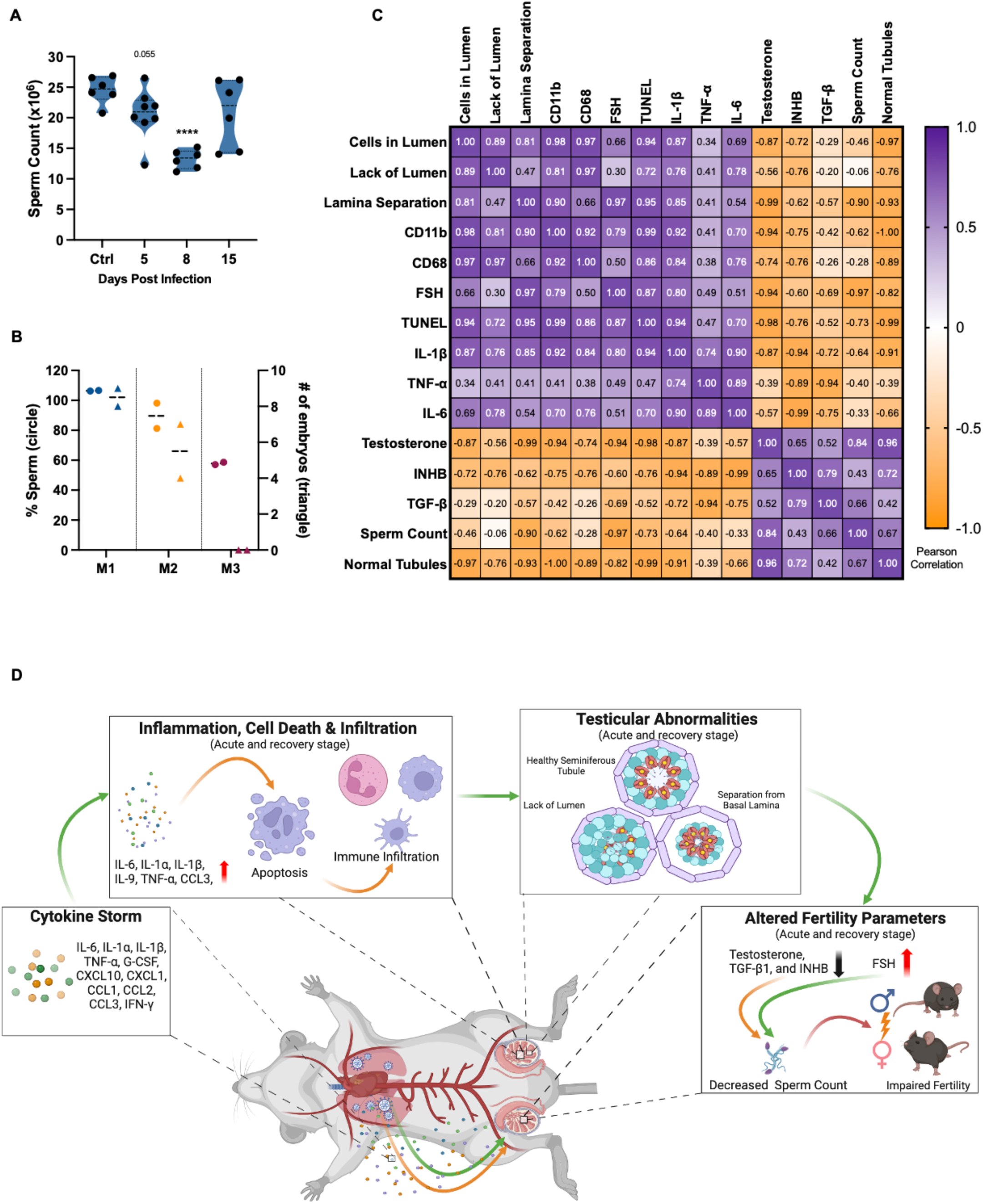
The epididymal sperm count declines during the acute phase of the disease and correlates with infertility in a subset of recovered mice. **(A)** Sperm count per epididymis (n, 3-5 mice per group) was measured at indicated time points. Each data points represents data from one epididymis. Significance (****p<0.0001) was determined using student’s t-test. **(B)** Relationship between percent sperm/epididymis (left y-axis) from each survivor male (circle, M1, blue; M2, yellow; M3, maroon) and number of fetuses (right y-axis, triangle) in each female (n, 2 females for each male). **(C)** Pearson’s correlation coefficient analysis of testicular defects, pro-inflammatory markers, cell death markers, and fertility parameters during the acute stage of the infection (D3 and D5) and early in the recovery stage (D8). **(D)** Schematic illustration of testicular injury proposed during SARS-CoV-2 infection.

The Pearson’s coefficient correlation matrix analysis of different injury and function markers clearly revealed that the testicular defects, immune cell infiltrates, cell death, and testicular cytokines have a strong positive correlation with each other (Fig. 8C). In contrast, a strong inverse relationship was observed between the injury markers and testis function markers including testosterone, inhibin B, sperm count, and TGF-β, indicating the profound effect SARS-CoV-2 exerts on male reproductive health (Fig. 8C).

## Discussion

The long-term effects of SARS-CoV-2 infection on male reproductive health including global changes at the molecular level and the extent of injury in recovered patients have not yet been characterized. We here extend our previous study to understand the testicular impairments, one of the symptoms of long COVID, during the acute stage of disease and in the survivor K18 hACE2 mice. We demonstrate that (i) the SARS-CoV-2 infection leads to defects of the seminiferous epithelium that resolve within one spermatogenesis cycle, (ii) testicular injury is associated with dysregulation of key biological pathways involved in spermatogenesis, BTB integrity, and immune homeostasis, (iii) the SARS-CoV-2-induced cytokine storm and testicular inflammation persists after virus clearance from the lung and is associated with cell death and immune cell infiltration in the testis, (iv) the hormonal imbalance is sustained after infection, especially testosterone levels, and correlates with transient reduction in the sperm count and the ability of survivor males to reproduce.

Although small animal models to study long COVID continue to be developed, there is currently no mouse model that recapitulates all aspects of PASC. Therefore, different models representing mild or severe disease are needed to study the pathophysiology of each symptom. Recently, a chronic SARS-CoV-2 infection model was reported that used an adeno-associated virus to deliver human ACE2 to the lungs of humanized MISTRG6 mice (39). This model exhibited persistent virus replication and chronic inflammation for 30 days in the lungs and represented rare cases in which persistent virus infection is seen in immunosuppressed patients (40). Infection of wild-type mice using SARS-CoV-2 mouse-adapted strain that has acquired several mutations not seen in human variants results in much milder disease symptoms (41, 42) and testicular injury has not been reported yet in this model. In our K18-hACE2 mice model, although survival is only 15-20%, the survivor mice represent moderate to severe disease in patients who are at a higher risk of developing PASC symptoms (4). Similar to what is seen with the post-mortem testicular tissue from COVID-19 patients (7, 9), infected mice during the acute stage of the disease display several pathogenic features in the seminiferous tubules. However, the long-term impact on male reproductive health in recovered COVID patients is only assessed based on fertility markers in the serum and/or semen. Our data extends the human histopathological observations to include different analyses of testes defects during the recovery stage and shows that severe tubular defects including sloughing of healthy spermatids and spermatocytes into the lumen and apoptotic cells persist after the acute stage of infection. Interestingly, separation of germinal epithelium from basal lamina and lack of tubular lumen are characteristic injury markers also seen in autoimmune orchitis and testicular cancer (43, 44).

Although, some of the morphological defects in the testis were resolved by D14, and all by D30, our RNA-seq analysis revealed pathways that were deregulated during peak disease and remained deregulated for longer time, including at D30. Several of these pathways were found associated with long-term testicular impairments. The downregulation of important pathways related to testicular architecture and function including focal adhesion kinase (FAK) and integrin-linked kinase (ILK) is a critical link to tubular defects and apoptosis. FAK regulates cell adhesion, migration, and survival, which are essential for the proper development and maturation of germ cells within the seminiferous tubules, and the integrity of the BTB (25, 45). FAK signaling is also tightly involved in tissue repair (46, 47) and its downregulation thereby suggests a potential delay in the repair of seminiferous tubules.

Persistent inflammation is a hallmark of long COVID patients (48). Our RNA Seq findings show that multiple inflammatory response pathways are significantly upregulated in the testis post SARS-CoV-2 infection, and that this correlate with significant increase in several pathogenic cytokines and chemokines for up to 4 weeks after infection. IL-12 signaling plays an important role in shifting the anti-inflammatory M2 macrophages to pro-inflammatory M1 and is shown to be activated at D8, D14, and D30 time points. Testicular M2 macrophages regulate spermatogenesis (49), and participate in the clearance of apoptotic cells and debris (50). Therefore, macrophage polarization to a pro-inflammatory state and tissue infiltration may usurp the immune-privileged environment of the testis. Increased testicular cytokines also correlate well with upregulation of the pathways associated with apoptosis, necrosis, fibrosis, and cellular infiltration (Fig. 2D). These data agrees with findings of immune cell infiltration and cell death in human post-mortem tissues (7, 9).

Many of the inflammatory cytokines in the testis overlapped with the cytokines induced by SARS-CoV-2 in other tissues and serum like IL-1β, IL-6 and TNF-α. However, IL-13, IL-9, and IL-10 were only induced in the testis. Elevated cytokines are observed in different models of testicular infection and cancer including mumps virus (MuV) that, unlike SARS-CoV-2, establishes productive infection in Sertoli and Leydig cells (51, 52). IL-6 has been shown to induce testicular germ cell death in experimental immune orchitis model in rats (53) while TNF-α has been shown to affect tight junction formation and integrity of the BTB (54, 55). We speculate that increased IL-6 and TNF-α in the testis after SARS-CoV-2 infection may directly impact the BTB integrity as also evidenced by the downregulation of critical pathways such as FAK and ILK signaling and decreased ZO-1 staining. The elevated levels of chemoattractant CXCL1 observed in our data is interesting and correlate with increased CD11b+ and CD68+ cells infiltration in the testis.

An interesting finding in our study is a significantly higher number of TUNEL-positive cells at D14 that remain relatively higher than controls even at D30. When compared to the early time points where most of the TUNEL-positive cells were found in the periphery of the seminiferous tubules, at D14 and D30, majority of TUNEL-positive cells appeared to be closer to the lumen of seminiferous tubules, indicative of more advanced germ cells being affected. We speculate one of the reasons for this might be delayed clearance of apoptotic cells due to reduced infiltrated immune cells and restoration of BTB integrity at D14 and D30. The trend of continued presence of apoptotic cells at D30 suggests that complete recovery may take longer than one spermatogenesis cycle, which is 34 days in mice and 74 days in humans.

In our data, the alterations in the important parameters associated with fertility were evident with a significant reduction in the serum testosterone levels lasting to D30. These data, along with the dysregulation in the steroidogenic pathway, suggest a long-term impact on Leydig cell function and aligns well with clinical data showing decreased testosterone levels in COVID-19 patients for up to 7 months after recovery (18, 56, 57). Increased FSH levels are also indicative of abnormal spermatogenesis and may suggest primary testicular failure due to its role in the feedback mechanism via the HPG axis following SARS-CoV-2 infection (58).

Another important highlight of our data is the finding of a significant reduction in the testicular TGF-β1 levels. Increased TGF-β1 has been shown in the serum from COVID-19 patients during the acute stage of the disease (59) and is proposed to be an important mediator of lung fibrosis (60). However, TGF-β1 is known to regulate multiple physiological functions in the testis, including spermatogenesis, Leydig cell steroidogenesis and extracellular matrix synthesis. The absence or overexpression of TGF-β1 has been shown to affect male reproductive function, supporting the fact that this signaling pathway is carefully coordinated to maintain testicular homeostasis and spermatogenesis (61). While our serum data agreed with other studies (62), decreased testicular levels of TGF-β1 is a new finding and suggests that it may be one of the underlying mechanisms involved in the dysfunction of testis function in recovering COVID-19 patients.

The assessment of the mid- and long-term impact of COVID-19 in humans revealed that decreased sperm quality and total sperm number improves after a recovery time of about 5 months (63), i.e. after almost two spermatogenesis cycle. Unlike humans, the decrease in the epididymal sperm count in mice was transient and recovered in two-third of mice within the first spermatogenesis cycle (D15). It is not clear whether decreased sperm count impacts male fertility in recovering long COVID patients. Although the limitation of our study is the small sample size for fertility testing and needs further validation, the results nevertheless indicate that the mice that do not efficiently recover the sperm count have a higher chance of transient infertility. This information is important clinically and may inform clinicians to consider medical interventions, if necessary, within about two months after recovery, to improve the fertility of male patients as soon as possible.

In summary, we evaluated several markers of male reproductive health and provided evidence that SARS-CoV-2 induces testicular injury and global alterations in the key biological pathways that mostly resolve within one spermatogenesis cycle in K18-hACE2 mice. As shown in the hypothetical model in Fig. 8D, we propose that SARS-CoV-2 acute lung infection triggers immune hyperactivation/dysregulation and excessive secretion of pro-inflammatory cytokines/chemokines in the systemic circulation in these mice. The cytokine storm induces a local inflammation in the testis, which alters the immune privileged environment of the seminiferous tubules and induces gross morphological alterations, apoptosis, disruption of the BTB, and recruitment of peripheral myeloid cells. These events collectively manifest in the deregulation of several fertility markers, many of which persist during the recovery phase. Our data also support that the hACE2 mouse permits the study of the mechanisms of long-term testicular injury, provides new insights beyond our current understanding of the impact of infection on male fertility, and validation of biomarkers to monitor the recovery of reproductive health. These findings suggest that recovering COVID-19 patients should be closely monitored to timely rescue the pathophysiological effects on male reproductive health.

## Materials and Methods

### Infection of *K18-hACE2* mice and virus quantitation

*B6.Cg-Tg(K18-ACE2)2Prlmn/J* (*K18-hACE2*) mice were obtained from the Jackson Laboratory and infected intranasally with 2×10^4^ PFU of SARS-CoV-2 USA-HI-B.1.429 isolated from a local COVID-19 patient that is very similar to the SARS-CoV-2 CoV/USA-WA1/2020 (64). All mouse experiments were performed on eight to twelve weeks old males at the dedicated ABSL3 facility according to the animal experimental guidelines issued and approved by the Institutional Animal Care and Use Committee of the University of Hawaii at Manoa. All mice were perfused intracardially with at least 10mL of PBS and lung, heart, and testis were either flash-frozen or fixed in 4% PFA and/or Bouin for various assays. At least 5 independent sets of experiments were conducted to obtain tissues from at least 3 survivor mice at all time points. Approximately 100-200μL of blood was collected via the tail vein, in Microvette® CB 300 Blood Collection System (Kent Scientific) and was centrifuged for 7 mins at 1000g. Serum was aliquoted and stored at -80 degrees Celsius. RNA was extracted from frozen tissues using QIAshredder and RNeasy Mini Plus Kit (Qiagen) and protein tissue lysates were prepared tissue protein extraction reagent (Thermo Fisher Sci. Cat. # FNN0071). SARS-CoV-2 titers in the normalized tissue homogenates were measured by plaque assay using Vero E6 cells as described previously (21). Intracellular viral genome copies were measured in the RNA extracted from cell lysates and tissue homogenates at different time points post-infection by qRT-PCR as described by us previously (21).

### Testis histology analysis

Testes were dissected and divided for different analyses. For histology, either full testes or halved testes were fixed in Bouin solution overnight and then stored in 70% ethanol prior to embedding in paraffin wax. Testis sections were cut at 5 μm thickness and stained with Periodic-acid Schiff and hematoxylin (PAS-H). The stages of seminiferous tubules were identified based on the composition of cells near the basal membrane according to the method described by Ahmed and de Rooij and as described by us before (65, 66). For the analysis of seminiferous epithelium abnormalities, 50 tubules from each male were analyzed and classified as normal or abnormal in regard to their organizational and germ cell features. Individual histopathological alterations including cells in lumen, lack of lumen, separation from the basal lamina, degenerating nucleus, and apoptotic cells were determined as described previously (65). To eliminate sample-to-sample variations arising from fixing of whole or halved testis, the statistical analyses of infected testis fixed as a whole was done by comparing to the control testis fixed similarly, and infected testis fixed after halving was compared to the control testis that was also halved before fixation.

### RNA Sequencing Analysis

RNA samples extracted from the whole testis from control and infected mice (n=3) at D3, 5, 8, 14, and 30 were used for transcriptomics assay. RNA integrity was measured using Agilent Bioanalyzer and sent to Medgenome Inc. for sequencing using Illumina TruSeq stranded mRNA kit (sequencer: NovaSeq). Alignment was performed using STAR (v2.7.3a) aligner. The raw read counts were estimated using HTSeq (v0.11.2). Read counts were normalized using DESeq2 to get the normalized counts. Additionally, the aligned reads were used for estimating the expression of the genes using cufflinks (v2.2.1). Principal component analysis (PCA) was performed for normalized counts of all the protein-coding genes from all the samples. Differential expression analysis was performed using DESeq2. The number of significantly up or down differentially expressed protein-coding genes without fold change shrinkage in the treated group (n=3/timepoint) vs. control group (n=3) were used for the analyses (Significant differential expression cutoff: P value < 0.05 & log2 fold-change >|1|). In addition, the low-quality sequence reads are excluded from the analysis. Data quality check was performed using FastQC (v0.11.8). The adapter trimming was performed using fastq-mcf (v1.05) and cutadapt (v2.5). Unwanted sequences include: mitochondrial genome sequences, ribosomal RNAs, transfer RNAs, adapter sequences and others. Contamination removal was performed using Bowtie2 (v2.5.1). Gene Ontology Over Representation Analysis (ORA) was done using enrichGO method from clusterProfiler (OrgDb/ AnnotationDbi: org.Mm.eg.dbv3.7.0 for Mouse from Bioconductor). Ingenuity Pathway Analysis (IPA) was used to determine the enrichment of biological functions and networks. IPA produced functional networks, pathways, and predicted upstream transcriptional regulators. IPA cutoff criteria for the input list of differentially expressed genes was set to log2 fold-change >|1| and p-value of <0.05 as described previously (67).

### Immunofluorescence and TUNEL assays

PFA-fixed tissues from mock and infected K18-hACE2 mouse testes were processed in sucrose and embedded in OCT. Sections (10μm thickness) cut using Leica UC7 microtome were permeabilized with 0.1% Triton X-100 in PBS and blocked with 5% bovine serum albumin in PBS. Sections were then incubated with primary antibodies against anti-CD68 (Invitrogen Cat. # 14-0681-82 at 1:250 dilution) and anti-CD11b (Invitrogen Cat. # 14-0112-82 at 1:250 dilution), followed by fluorophore-conjugated secondary antibody (Invitrogen Alexa Fluor 594-conjugated, at 1:1000 dilution), and examined using a Leica SP3, DMI8 confocal microscope for CD68 and CD11b images. TUNEL assay was performed using the Invitrogen Click-iT™ Plus TUNEL Assay Kit (Thermo Fisher Sci Cat. # C10617) according to the manufacturer’s instructions. ImageJ/Fiji software was used for densitometry of all the representative images using the average of 3 different fields from 3 different mice per group.

### Multi-plex and single-plex Enzyme-Linked Immunosorbent Assays

Lung, heart, and testis tissue lysates were prepared in tissue extraction reagent (Thermo Fisher Sci Cat. # FNN0071) with 1X protease and phosphatase inhibitor (Thermo Fisher Sci Cat. # 78440) by using a sonicator. Kits used for ELISA were TGF-β1 Mouse ELISA Kit (Thermo Fisher Sci Cat. # BMS608-4TWO), Mouse Testosterone ELISA Kit (Crystal Chem. Cat. # 80552), Mouse FSH ELISA Kit (Abclonal Cat. # RK04237), Mouse INHB ELISA kit (Abclonal Cat. # RK02949), Mouse LH ELISA Kit (Abclonal Cat. # RK02986). All assays using normalized protein amounts were performed according to the manufacturer’s instructions. MILLIPLEX Mouse Cytokine/Chemokine Magnetic Bead Panel (Millipore Sigma Cat. # MCYTOMAG-70K-PMX) was used to measure 27 cytokines and chemokines as described previously (68).

### Epididymal sperm count and fertility assays

Cauda epidydimal sperm from each mouse were released in 1 mL of PBS and incubated for 30 min at 37°C as described previously (69). Spermatozoa were extracted by filtering through 70μm filters, and the total sperm count was assessed using hemacytometer. Fertility was tested for 3 recovered males who were proven fertile before infection. At D10 post infection, each recovered male was placed in a new cage with two females for 6 days. At D15, males were sacrificed to measure sperm count, and the females were sacrificed after 10 days to assess implantation sites reflective of embryo count.

### Statistical Analysis

SARS-CoV-2 titers are reported as means +/- standard error of the mean (SEM) of data from ≥3 independent experiments. Statistically significant difference between data from different groups was determined by unpaired Student’s t-test or Mann-Whitney test using GraphPad Prism 10.0.2 (GraphPad Software, San Diego, CA). Simple linear regression analysis, Pearson’s correlations, and matrices were performed using GraphPad Prism software. A P value of <0.05 and a Pearson correlation coefficient >|0.6| were considered statistically significant for all analyses.

---

**Figure S1.**
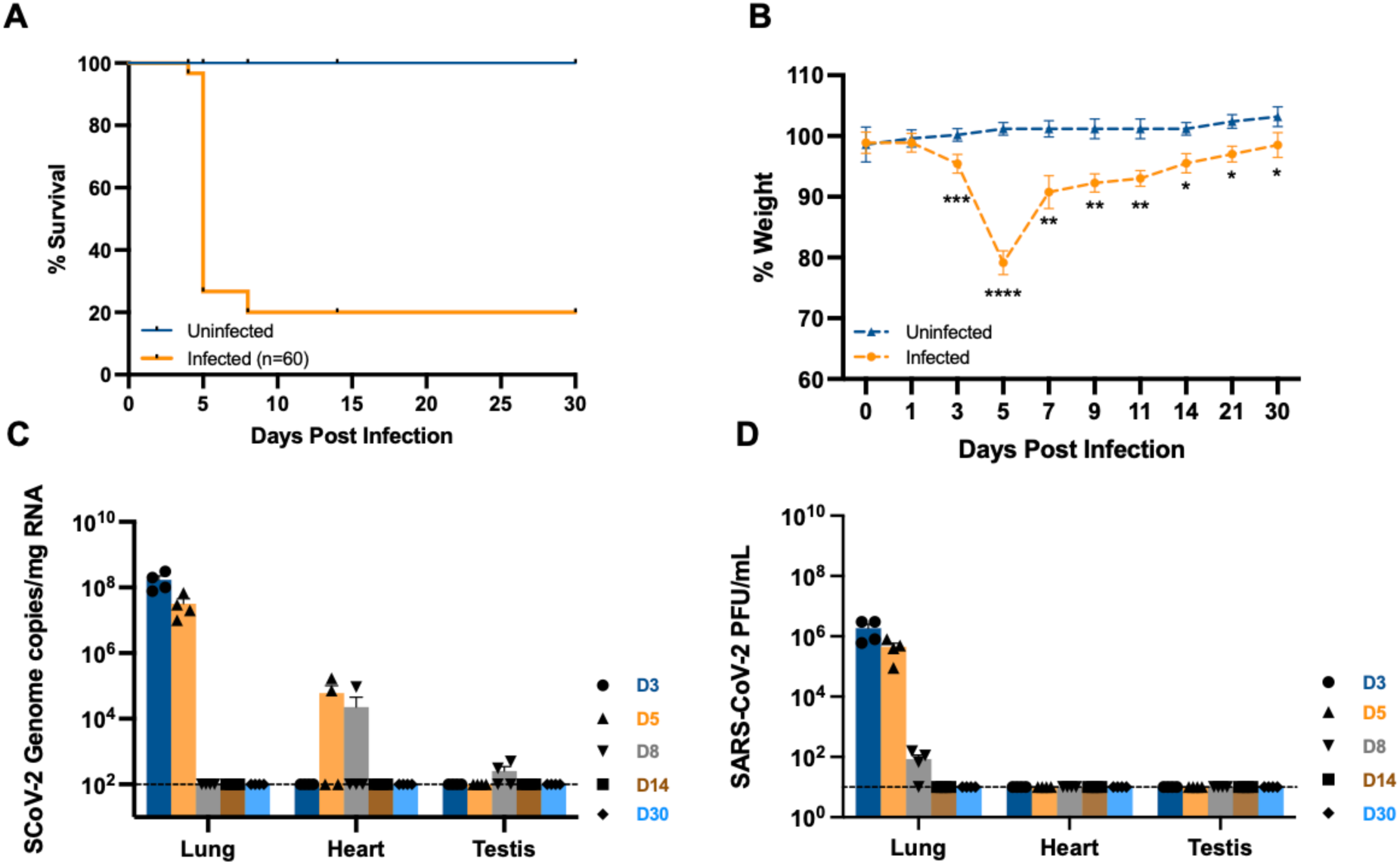
SARS-CoV-2 infection of K18-hACE2 mice. **(A)** Survival curve, and **(B)** Percent weight change of intranasally infected K18-hACE2 mice at indicated time points. **(C)** SARS-CoV-2 genome copies in the lung, heart, brain, and testis at indicated time points measured using qRT-PCR and expressed as copies/mg RNA. **(D)** Plaque assay of progeny SARS-CoV-2 titers in the tissue homogenates of lung, heart, testis, and brain measured at indicated time points. Significance *p<0.05, **p<0.01, ***p<0.001, ****p<0.0001 was determined using student’s t-test.

**Figure S2.**
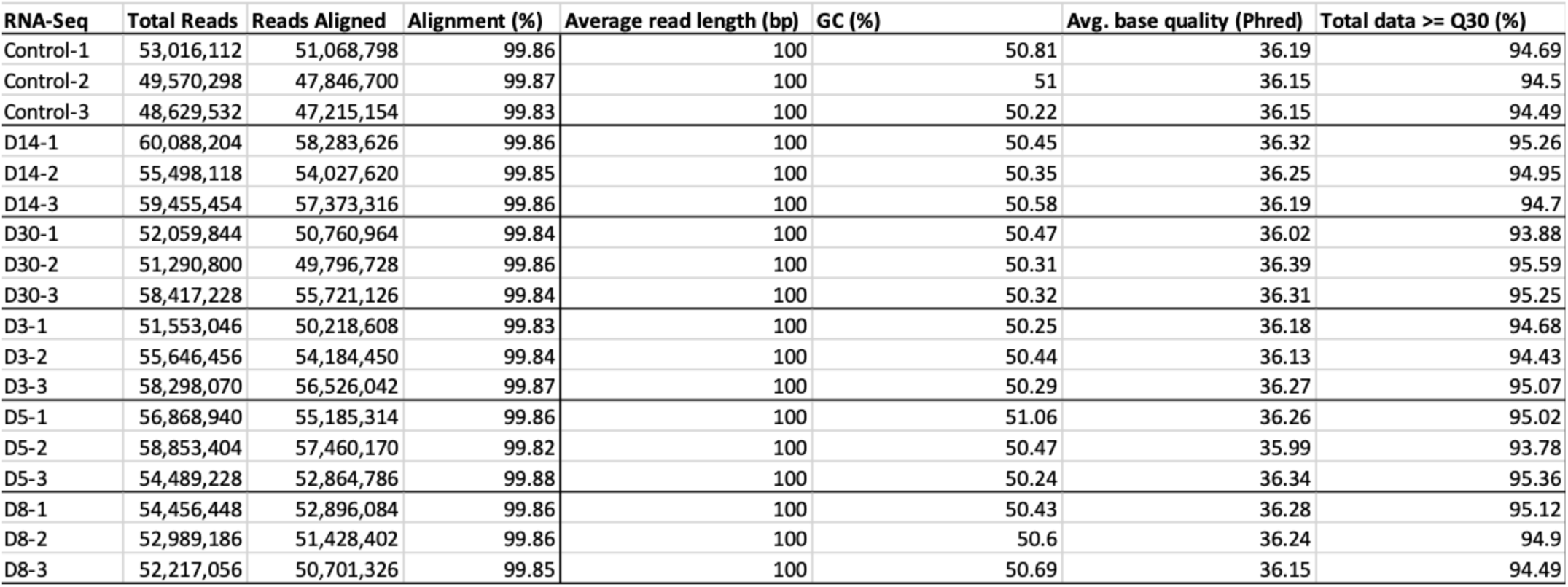
Quality Control of RNA samples used for RNA-Seq Analysis. : total reads, aligned reads, percent alignment, average read length, percent GC, Phred score, and percent Q30 of all the samples submitted for RNA-seq analysis (n, 3/timepoint D3, 5, 8, 14, 30).

**Figure S3.**
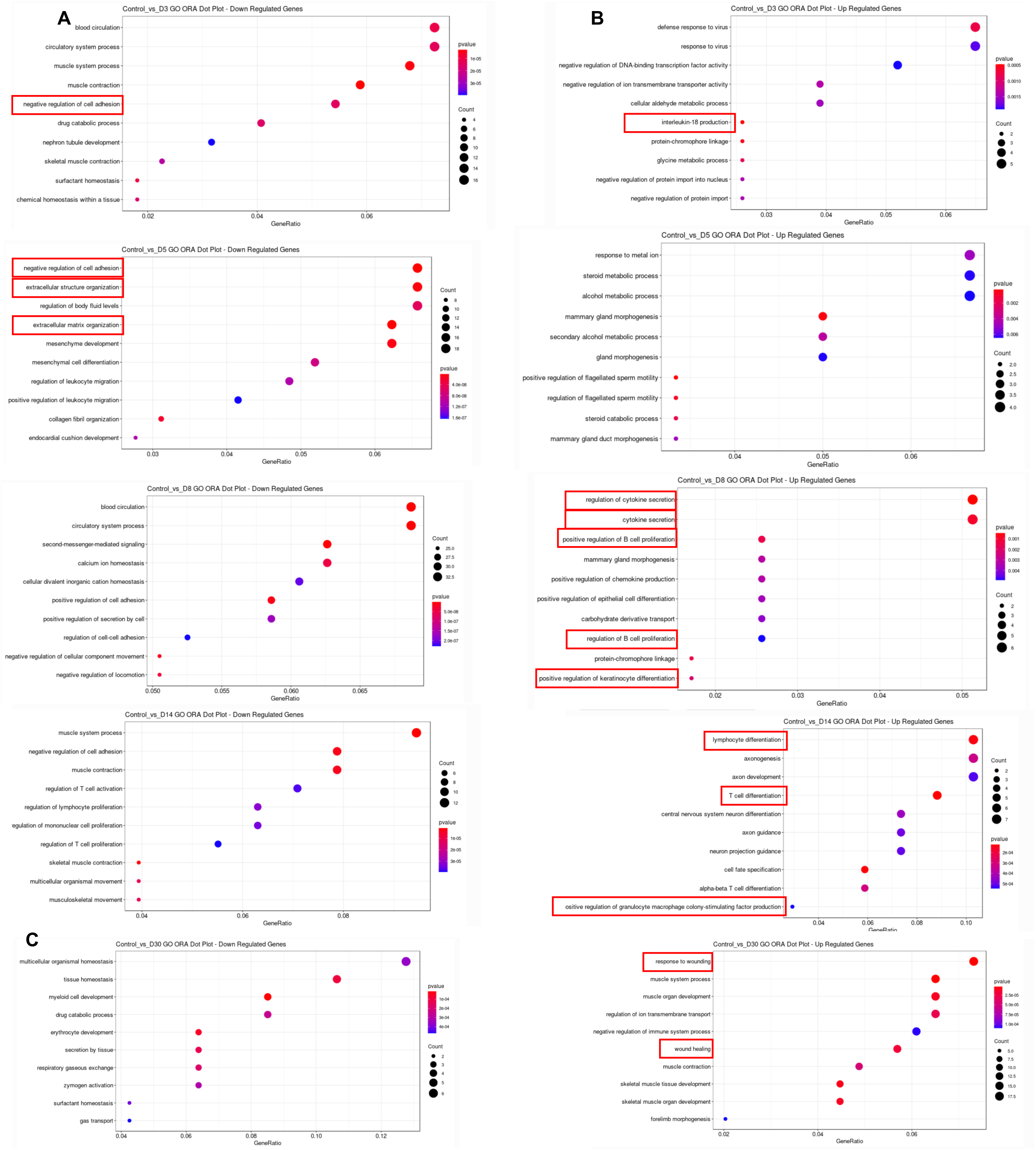
Gene ontology (GO) overrepresentation analysis (ORA) of testis from SARS-CoV-2-infected mice. **(A)** Top downregulated gene pathways were identified at D3, D5, D8 and D14 with respect to control. **(B)** Top upregulated gene pathways observed at D3, D5, D8 and D14 with respect to control. (C) Top upregulated gene pathways at D30 (right) and downregulated gene pathways at D30 (left) with respect to control. Cutoffs for all analysis was p<0.05 and log2fc ≥ |1|

## Notes

### Competing Interest Statement

The authors have declared no competing interest.

## References

1. Al-Aly Z, Xie Y, Bowe B. 2021. High-dimensional characterization of post-acute sequelae of COVID-19. Nature 594:259–264.

2. Chen C, Haupert SR, Zimmermann L, Shi X, Fritsche LG, Mukherjee B. 2022. Global Prevalence of Post-Coronavirus Disease 2019 (COVID-19) Condition or Long COVID: A Meta-Analysis and Systematic Review. J Infect Dis 226:1593–1607.

3. Lopez-Leon S, Wegman-Ostrosky T, Ayuzo Del Valle NC, Perelman C, Sepulveda R, Rebolledo PA, Cuapio A, Villapol S. 2022. Long-COVID in children and adolescents: a systematic review and meta-analyses. Sci Rep 12:9950.

4. Davis HE, McCorkell L, Vogel JM, Topol EJ. 2023. Long COVID: major findings, mechanisms and recommendations. Nat Rev Microbiol 21:133–146.

5. Subramanian A, Nirantharakumar K, Hughes S, Myles P, Williams T, Gokhale KM, Taverner T, Chandan JS, Brown K, Simms-Williams N, Shah AD, Singh M, Kidy F, Okoth K, Hotham R, Bashir N, Cockburn N, Lee SI, Turner GM, Gkoutos GV, Aiyegbusi OL, McMullan C, Denniston AK, Sapey E, Lord JM, Wraith DC, Leggett E, Iles C, Marshall T, Price MJ, Marwaha S, Davies EH, Jackson LJ, Matthews KL, Camaradou J, Calvert M, Haroon S. 2022. Symptoms and risk factors for long COVID in non-hospitalized adults. Nat Med 28:1706–1714.

6. Maleki BH, Tartibian B. 2021. COVID-19 and male reproductive function: a prospective, longitudinal cohort study. Reproduction 161:319–331.

7. Duarte-Neto AN, Teixeira TA, Caldini EG, Kanamura CT, Gomes-Gouvêa MS, Dos Santos ABG, Monteiro RAA, Pinho JRR, Mauad T, da Silva LFF, Saldiva PHN, Dolhnikoff M, Leite KRM, Hallak J. 2022. Testicular pathology in fatal COVID-19: A descriptive autopsy study. Andrology 10:13–23.

8. Yang M, Chen S, Huang B, Zhong J-M, Su H, Chen Y-J, Cao Q, Ma L, He J, Li X- F, Li X, Zhou J-J, Fan J, Luo D-J, Chang X-N, Arkun K, Zhou M, Nie X. 2020. Pathological Findings in the Testes of COVID-19 Patients: Clinical Implications. Eur Urol Focus 6:1124–1129.

9. Ma X, Guan C, Chen R, Wang Y, Feng S, Wang R, Qu G, Zhao S, Wang F, Wang X, Zhang D, Liu L, Liao A, Yuan S. 2021. Pathological and molecular examinations of postmortem testis biopsies reveal SARS-CoV-2 infection in the testis and spermatogenesis damage in COVID-19 patients. 2. Cell Mol Immunol 18:487–489.

10. Gacci M, Coppi M, Baldi E, Sebastianelli A, Zaccaro C, Morselli S, Pecoraro A, Manera A, Nicoletti R, Liaci A, Bisegna C, Gemma L, Giancane S, Pollini S, Antonelli A, Lagi F, Marchiani S, Dabizzi S, Degl’Innocenti S, Annunziato F, Maggi M, Vignozzi L, Bartoloni A, Rossolini GM, Serni S. 2021. Semen impairment and occurrence of SARS-CoV-2 virus in semen after recovery from COVID-19. Human Reproduction 36:1520–1529.

11. Schroeder M, Schaumburg B, Mueller Z, Parplys A, Jarczak D, Roedl K, Nierhaus A, de Heer G, Grensemann J, Schneider B, Stoll F, Bai T, Jacobsen H, Zickler M, Stanelle-Bertram S, Klaetschke K, Renne T, Meinhardt A, Aberle J, Hiller J, Peine S, Kreienbrock L, Klingel K, Kluge S, Gabriel G. 2021. Sex hormone dysregulations are associated with disease severity in critically ill male COVID-19 patients - a retrospective analysi 10.21203/rs.3.rs-476932/v1.

12. Rastrelli G, Di Stasi V, Inglese F, Beccaria M, Garuti M, Di Costanzo D, Spreafico F, Greco GF, Cervi G, Pecoriello A, Magini A, Todisco T, Cipriani S, Maseroli E, Corona G, Salonia A, Lenzi A, Maggi M, De Donno G, Vignozzi L. 2021. Low testosterone levels predict clinical adverse outcomes in SARS-CoV-2 pneumonia patients. Andrology 9:88–98.

13. Guo L, Zhao S, Li W, Wang Y, Li L, Jiang S, Ren W, Yuan Q, Zhang F, Kong F, Lei J, Yuan M. 2021. Absence of SARS-CoV-2 in semen of a COVID-19 patient cohort. Andrology 9:42–47.

14. Holtmann N, Edimiris P, Andree M, Doehmen C, Baston-Buest D, Adams O, Kruessel J-S, Bielfeld AP. 2020. Assessment of SARS-CoV-2 in human semen-a cohort study. Fertil Steril 114:233–238.

15. Li D, Jin M, Bao P, Zhao W, Zhang S. 2020. Clinical Characteristics and Results of Semen Tests Among Men With Coronavirus Disease 2019. JAMA Network Open 3:e208292.

16. Martinez MS, Ferreyra FN, Paira DA, Rivero VE, Olmedo JJ, Tissera AD, Molina RI, Motrich RD. 2023. COVID-19 associates with semen inflammation and sperm quality impairment that reverses in the short term after disease recovery. Front Physiol 14:1220048.

17. Salonia A, Pontillo M, Capogrosso P, Pozzi E, Ferrara AM, Cotelessa A, Belladelli F, Corsini C, Gregori S, Rowe I, Carenzi C, Ramirez GA, Tresoldi C, Locatelli M, Cavalli G, Dagna L, Castagna A, Zangrillo A, Tresoldi M, Landoni G, Rovere-Querini P, Ciceri F, Montorsi F. 2023. Testosterone in males with COVID-19: a 12-month cohort study. Andrology 11:17–23.

18. Salonia A, Pontillo M, Capogrosso P, Gregori S, Carenzi C, Ferrara AM, Rowe I, Boeri L, Larcher A, Ramirez GA, Tresoldi C, Locatelli M, Cavalli G, Dagna L, Castagna A, Zangrillo A, Tresoldi M, Landoni G, Rovere-Querini P, Ciceri F, Montorsi F. 2022. Testosterone in males with COVID-19: A 7-month cohort study. Andrology 10:34–41.

19. Zhao S, Zhu W, Xue S, Han D. 2014. Testicular defense systems: immune privilege and innate immunity. Cell Mol Immunol 11:428–437.

20. Le Tortorec A, Matusali G, Mahé D, Aubry F, Mazaud-Guittot S, Houzet L, Dejucq-Rainsford N. 2020. From Ancient to Emerging Infections: The Odyssey of Viruses in the Male Genital Tract. Physiol Rev 100:1349–1414.

21. Giannakopoulos S, Strange DP, Jiyarom B, Abdelaal O, Bradshaw AW, Nerurkar VR, Ward MA, Bakse J, Yap J, Vanapruks S, Boisvert WA, Tallquist MD, Shikuma C, Sadri-Ardekani H, Clapp P, Murphy S, Verma S. 2023. In vitro evidence against productive SARS-CoV-2 infection of human testicular cells: Bystander effects of infection mediate testicular injury. PLOS Pathogens 19:e1011409.

22. Bhaskar S, Sinha A, Banach M, Mittoo S, Weissert R, Kass JS, Rajagopal S, Pai AR, Kutty S. 2020. Cytokine Storm in COVID-19—Immunopathological Mechanisms, Clinical Considerations, and Therapeutic Approaches: The REPROGRAM Consortium Position Paper. Frontiers in Immunology 11.

23. Mustafa MI, Abdelmoneim AH, Mahmoud EM, Makhawi AM. 2020. Cytokine Storm in COVID-19 Patients, Its Impact on Organs and Potential Treatment by QTY Code-Designed Detergent-Free Chemokine Receptors. Mediators Inflamm 2020:8198963.

24. Schultheiß C, Willscher E, Paschold L, Gottschick C, Klee B, Henkes S-S, Bosurgi L, Dutzmann J, Sedding D, Frese T, Girndt M, Höll JI, Gekle M, Mikolajczyk R, Binder M. 2022. The IL-1β, IL-6, and TNF cytokine triad is associated with post-acute sequelae of COVID-19. Cell Rep Med 3:100663.

25. Siu ER, Wong EWP, Mruk DD, Porto CS, Cheng CY. 2009. Focal adhesion kinase is a blood–testis barrier regulator. Proceedings of the National Academy of Sciences 106:9298–9303.

26. Yen C-F, Wang H-S, Lee C-L, Liao S-K. 2014. Roles of integrin-linked kinase in cell signaling and its perspectives as a therapeutic target. Gynecology and Minimally Invasive Therapy 3:67–72.

27. Laing NG, Dye DE, Wallgren-Pettersson C, Richard G, Monnier N, Lillis S, Winder TL, Lochmüller H, Graziano C, Mitrani-Rosenbaum S, Tuomey D, Sparrow JC, Beggs AH, Nowak KJ. 2009. Mutations and Polymorphisms of the Skeletal Muscle α-Actin Gene (ACTA1). Hum Mutat 30:1267–1277.

28. Richards CD. 2013. The Enigmatic Cytokine Oncostatin M and Roles in Disease. ISRN Inflamm 2013:512103.

29. Wang Y, Xie L, Tian E, Li X, Wen Z, Li L, Chen L, Zhong Y, Ge R. 2019. Oncostatin M inhibits differentiation of rat stem Leydig cells in vivo and in vitro. J Cell Mol Med 23:426–438.

30. Broxmeyer HE, Cooper SH, Ropa J. 2021. CXCL15/Lungkine has suppressive activity on proliferation and expansion of multi-potential, erythroid, granulocyte and macrophage progenitors in S-phase specific manner. Blood Cells Mol Dis 91:102594.

31. Gianzo M, Subirán N. 2020. Regulation of Male Fertility by the Renin-Angiotensin System. Int J Mol Sci 21:7943.

32. Wang J, Li J, Xu W, Xia Q, Gu Y, Song W, Zhang X, Yang Y, Wang W, Li H, Zou K. 2019. Androgen promotes differentiation of PLZF+ spermatogonia pool via indirect regulatory pattern. Cell Communication and Signaling 17:57.

33. Lindeboom F, Gillemans N, Karis A, Jaegle M, Meijer D, Grosveld F, Philipsen S. 2003. A tissue-specific knockout reveals that Gata1 is not essential for Sertoli cell function in the mouse. Nucleic Acids Res 31:5405–5412.

34. Viger RS, de Mattos K, Tremblay JJ. 2022. Insights Into the Roles of GATA Factors in Mammalian Testis Development and the Control of Fetal Testis Gene Expression. Front Endocrinol (Lausanne) 13:902198.

35. Abudureheman T, Xia J, Li M-H, Zhou H, Zheng W-W, Zhou N, Shi R-Y, Zhu J-M, Yang L-T, Chen L, Zheng L, Xue K, Qing K, Duan C-W. 2021. CDK7 Inhibitor THZ1 Induces the Cell Apoptosis of B-Cell Acute Lymphocytic Leukemia by Perturbing Cellular Metabolism. Front Oncol 11:663360.

36. Jankowska K, Suszczewicz N, Rabijewski M, Dudek P, Zgliczyński W, Maksym RB. 2022. Inhibin-B and FSH Are Good Indicators of Spermatogenesis but Not the Best Indicators of Fertility. Life (Basel) 12:511.

37. Gordetsky J, van Wijngaarden E, O’Brien J. 2012. Redefining abnormal follicle-stimulating hormone in the male infertility population. BJU Int 110:568–572.

38. Arguinchona LM, Zagona-Prizio C, Joyce ME, Chan ED, Maloney JP. 2023. Microvascular significance of TGF-β axis activation in COVID-19. Front Cardiovasc Med 9:1054690.

39. Sefik E, Israelow B, Mirza H, Zhao J, Qu R, Kaffe E, Song E, Halene S, Meffre E, Kluger Y, Nussenzweig M, Wilen CB, Iwasaki A, Flavell RA. 2022. A humanized mouse model of chronic COVID-19. 6. Nat Biotechnol 40:906–920.

40. Corey L, Beyrer C, Cohen MS, Michael NL, Bedford T, Rolland M. 2021. SARS-CoV-2 variants in immunosuppressed individuals. N Engl J Med 385:562–566.

41. Bader SM, Cooney JP, Sheerin D, Taiaroa G, Harty L, Davidson KC, Mackiewicz L, Dayton M, Wilcox S, Whitehead L, Rogers KL, Georgy SR, Coussens AK, Grimley SL, Corbin V, Pitt M, Coin L, Pickering R, Thomas M, Allison CC, McAuley J, Purcell DFJ, Doerflinger M, Pellegrini M. 2023. SARS-CoV-2 mouse adaptation selects virulence mutations that cause TNF-driven age-dependent severe disease with human correlates. Proceedings of the National Academy of Sciences 120:e2301689120.

42. Pan T, Chen R, He X, Yuan Y, Deng X, Li R, Yan H, Yan S, Liu J, Zhang Y, Zhang X, Yu F, Zhou M, Ke C, Ma X, Zhang H. 2021. Infection of wild-type mice by SARS-CoV-2 B.1.351 variant indicates a possible novel cross-species transmission route. 1. Sig Transduct Target Ther 6:1–12.

43. Rio R del, McAllister RD, Meeker ND, Wall EH, Bond JP, Kyttaris VC, Tsokos GC, Tung KSK, Teuscher C. 2012. Identification of Orch3, a Locus Controlling Dominant Resistance to Autoimmune Orchitis, as Kinesin Family Member 1C. PLOS Genetics 8:e1003140.

44. Stang A, McMaster ML, Sesterhenn IA, Rapley E, Huddart R, Heimdal K, McGlynn KA, Oosterhuis JW, Greene MH. 2021. Histological Features of Sporadic and Familial Testicular Germ Cell Tumors Compared and Analysis of Age-Related Changes of Histology. 7. Cancers 13:1652.

45. Li F, Zhang Y, Wu C. 1999. Integrin-linked kinase is localized to cell-matrix focal adhesions but not cell-cell adhesion sites and the focal adhesion localization of integrin-linked kinase is regulated by the PINCH-binding ANK repeats. Journal of Cell Science 112:4589–4599.

46. Januszyk M, Kwon SH, Wong VW, Padmanabhan J, Maan ZN, Whittam AJ, Major MR, Gurtner GC. 2017. The Role of Focal Adhesion Kinase in Keratinocyte Fibrogenic Gene Expression. Int J Mol Sci 18:1915.

47. Dai P, Qiao F, Chen Y, Chan DYL, Yim HCH, Fok KL, Chen H. 2023. SARS-CoV-2 and male infertility: from short- to long-term impacts. J Endocrinol Invest 1–17.

48. Talla A, Vasaikar SV, Szeto GL, Lemos MP, Czartoski JL, MacMillan H, Moodie Z, Cohen KW, Fleming LB, Thomson Z, Okada L, Becker LA, Coffey EM, De Rosa SC, Newell EW, Skene PJ, Li X, Bumol TF, Juliana McElrath M, Torgerson TR. 2023. Persistent serum protein signatures define an inflammatory subcategory of long COVID. 1. Nat Commun 14:3417.

49. DeFalco T, Potter SJ, Williams AV, Waller B, Kan MJ, Capel B. 2015. Macrophages Contribute to the Spermatogonial Niche in the Adult Testis. Cell Rep 12:1107–1119.

50. Washburn RL, Hibler T, Kaur G, Dufour JM. 2022. Sertoli Cell Immune Regulation: A Double-Edged Sword. Front Immunol 13:913502.

51. Wu H, Wang F, Tang D, Han D. 2021. Mumps Orchitis: Clinical Aspects and Mechanisms. Frontiers in Immunology 12.

52. Klein B, Haggeney T, Fietz D, Indumathy S, Loveland KL, Hedger M, Kliesch S, Weidner W, Bergmann M, Schuppe H-C. 2016. Specific immune cell and cytokine characteristics of human testicular germ cell neoplasia. Human Reproduction 31:2192–2202.

53. Rival C, Theas MS, Guazzone VA, Lustig L. 2006. Interleukin-6 and IL-6 receptor cell expression in testis of rats with autoimmune orchitis. J Reprod Immunol 70:43– 58.

54. Siu MKY, Lee WM, Cheng CY. 2003. The interplay of collagen IV, tumor necrosis factor-alpha, gelatinase B (matrix metalloprotease-9), and tissue inhibitor of metalloproteases-1 in the basal lamina regulates Sertoli cell-tight junction dynamics in the rat testis. Endocrinology 144:371–387.

55. Li MWM, Xia W, Mruk DD, Wang CQF, Yan HHN, Siu MKY, Lui W, Lee WM, Cheng CY. Tumor necrosis factor α reversibly disrupts the blood–testis barrier and impairs Sertoli–germ cell adhesion in the seminiferous epithelium of adult rat testes. Journal of Endocrinology 190:313–329.

56. Çayan S, Uğuz M, Saylam B, Akbay E. 2020. Effect of serum total testosterone and its relationship with other laboratory parameters on the prognosis of coronavirus disease 2019 (COVID-19) in SARS-CoV-2 infected male patients: a cohort study. Aging Male 23:1493–1503.

57. Okçelik S. 2021. COVID-19 pneumonia causes lower testosterone levels. Andrologia 53:e13909.

58. Gordetsky J, van Wijngaarden E, O’Brien J. 2012. Redefining abnormal follicle-stimulating hormone in the male infertility population. BJU Int 110:568–572.

59. Arguinchona LM, Zagona-Prizio C, Joyce ME, Chan ED, Maloney JP. 2023. Microvascular significance of TGF-β axis activation in COVID-19. Front Cardiovasc Med 9:1054690.

60. Vaz de Paula CB, Nagashima S, Liberalesso V, Collete M, da Silva FPG, Oricil AGG, Barbosa GS, da Silva GVC, Wiedmer DB, da Silva Dezidério F, Noronha L. 2021. COVID-19: Immunohistochemical Analysis of TGF-β Signaling Pathways in Pulmonary Fibrosis. Int J Mol Sci 23:168.

61. Itman C, Mendis S, Barakat B, Loveland KL. 2006. All in the family: TGF-beta family action in testis development. Reproduction 132:233–246.

62. Wang E-Y, Chen H, Sun B-Q, Wang H, Qu H-Q, Liu Y, Sun X-Z, Qu J, Fang Z-F, Tian L, Zeng Y-F, Huang S-K, Hakonarson H, Liu Z-G. 2021. Serum levels of the IgA isotype switch factor TGF-β1 are elevated in patients with COVID-19. FEBS Lett 595:1819–1824.

63. Hu B, Liu K, Ruan Y, Wei X, Wu Y, Feng H, Deng Z, Liu J, Wang T. 2022. Evaluation of mid- and long-term impact of COVID-19 on male fertility through evaluating semen parameters. Transl Androl Urol 11:159–167.

64. Maison DP, Ching LL, Cleveland SB, Tseng AC, Nakano E, Shikuma CM, Nerurkar VR. 2022. Dynamic SARS-CoV-2 emergence algorithm for rationally-designed logical next-generation vaccines. 1. Commun Biol 5:1–12.

65. Ruthig VA, Nielsen T, Riel JM, Yamauchi Y, Ortega EA, Salvador Q, Ward MA. 2017. Testicular abnormalities in mice with Y chromosome deficiencies. Biol Reprod 96:694–706.

66. Ahmed EA, de Rooij DG. 2009. Staging of mouse seminiferous tubule cross-sections. Methods Mol Biol 558:263–277.

67. Strange DP, Green R, Siemann DN, Gale M, Verma S. 2018. Immunoprofiles of human Sertoli cells infected with Zika virus reveals unique insights into host-pathogen crosstalk. Sci Rep 8:8702.

68. Siemann DN, Strange DP, Maharaj PN, Shi P-Y, Verma S. 2017. Zika Virus Infects Human Sertoli Cells and Modulates the Integrity of the In Vitro Blood-Testis Barrier Model. J Virol 91:e00623–17.

69. Holmlund H, Yamauchi Y, Durango G, Fujii W, Ward MA. 2022. Two acquired mouse Y chromosome-linked genes, Prssly and Teyorf1, are dispensable for male fertility‡. Biology of Reproduction 107:752–764.

